# Genomic Insights into the Evolution of Carnivory in the Giant Butterwort, *Pinguicula gigantea*: Chromosome-Level Assembly and Phylogenomic Analysis

**DOI:** 10.1101/2025.04.05.646448

**Authors:** Steven J. Fleck, Jonathan Kirshner, Sitaram Rajaraman, Tianying Lan, Crystal Tomlin, Luis Herrera-Estrella, Richard W. Jobson, Enrique Ibarra-Laclette, Martín Mata Rosas, Todd P. Michael, Victor A. Albert

## Abstract

Carnivorous plants provide a powerful system for studying plant morphological and physiological evolution. These plants capture and digest prey for nutrients and this rare trait has evolved independently at least eleven times in thirteen families within six orders. Among these are the butterworts in the genus *Pinguicula*. This group of plants captures prey on a basal rosette of sticky leaves where the prey is broken down and digested. Here we present the chromosome-level reference assembly of the carnivorous giant butterwort, *Pinguicula gigantea*. An additional two assemblies were generated for this study, primarily from long-read sequencing data. In this study, we focus on the genome evolution of *Pinguicula*, including confirming at least two whole genome duplications since the gamma hexaploidy event at the base of the core eudicots. We provide evidence from full-genome data that the carnivorous family that *Pinguicula* belongs to, the Lentibulariaceae, is most closely related to the Acanthaceae. We also report multiple tandem duplications of candidate digestive enzyme genes, including a putative tandem duplication of cysteine proteases that is syntenic with two cysteine protease tandem duplication in *Utricularia gibba* that contain genes with trap specific expression. We also show evidence that these tandem duplications evolved independently in both *Pinguicula* and *Utricularia gibbaUtricularia gibba* and were potentially facilitated by different repeat element families.

## Introduction

The carnivorous syndrome has evolved independently at least eleven times within thirteen families and six orders of flowering plants (reviewed in ^1–4^). Carnivorous plants have evolved traits that enable them to obtain most of their nutrients from animal sources and evolved complex strategies for capturing and digesting prey^5^. The Lentibulariaceae represents one origin of carnivorous plants and about 45% of all carnivorous plant species: ninety-five species (11%) in *Pinguicula*, thirty species (4%) in *Genlisea*, and 252 species (30%) in *Utricularia*. While this family has only a single proposed origin for carnivory, it has three divergent trapping strategies. *Utricularia* uses subterranean or aquatic bladder traps that actively suction in and capture prey, *Genlisea* use root-like “eel trap” structures with inward-facing hairs so prey can only enter and not exit, and *Pinguicula* leaves are covered with stalked adhesive-producing glands that passively capture prey^6^.

Several genomes from the Lentibulariaceae have been published. Genome assemblies and annotations have been made publicly available for *Genlisea aurea*^7^, *Utricularia gibba*^8^, and *Utricularia reniformis*^9^. Two additional genome assemblies and annotations were generated for *Genlisea hispidula* and *Genlisea nigrocaulis*, but these have not been made publicly available^10^. Lastly, genome assemblies have been published for *Pinguicula moranensis* and *Pinguicula primuliflora*^11^. Of these, only *Utricularia gibba* has some chromosome-level scaffolds (100,232,236 total length, 516 scaffolds, 8,502,017 N50, 10 L50). Four scaffolds have telomeric repeats at both ends and twenty scaffolds with telomeric repeats on one end^8^. The remaining Lentibulariaceae genome assemblies are much less contiguous. For example, *Pinguicula moranensis* has a total length of 816,101,522, made up of 255,161 contigs, and has an N50 of 5,325 and an L50 of 40,354. Assembly stats for all published carnivorous plant nuclear genomes can be find in Table S1.

Tandemly duplicated genes have been documented as being important to the evolution of the carnivorous syndrome. The genome assembly of *Utricularia gibba*, a family member and close relative of *Pinguicula*, contains tandem arrays of cysteine protease genes with trap specific expression^8^. Furthermore, significantly enriched gene ontology (GO) terms for tandem duplicates with trap-enhanced expression included peptide transport, ATPase activities, hydrolase and chitinase activities, and cell wall dynamic components potentially related to active bladder movements. All of these are related to trapping prey, trap acidification, prey breakdown, and intercellular movement of broken-down prey proteins^8^. Notably, the cysteine protease gene cluster is located within regions of LArge Retrotransposon Derivatives or LARDs^8^, suggesting their potential role in the duplication process.

*Nepenthes gracilis* also demonstrated the importance of tandem duplication in the evolution of its carnivorous syndrome^12^. Tandem clusters that encode for digestive enzymes had the highest mRNA abundance in the digestive zone of the pitchers and had increased expression following feeding. It was also found that *Nepenthes* had a decaploid structure, resulting in five haploid subgenomes with strong evidence of subgenome dominance. For each gene-rich dominant chromosome, there were four homologous gene-poor recessive chromosomes. Many trap-specific tandem clusters were located on recessive subgenomes, likely due to the relaxation of purifying selection on these chromosomes compared with the dominant ones, allowing for the potential sub- and neofunctionalization of digestive enzymes^12^. This highlights the utility of gene redundancy through whole genome duplication in the evolution of the carnivorous syndrome.

Other studies have used mass spectrometry to identify proteins in the digestive secretion of *Pinguicula*. The digestive secretions of *Pinguicula x tina*, a hybrid between *Pinguicula agnata* and *Pinguicula zecheri*, contained fourteen protein sequences covering eight different catalytic activities: three secretory proteases (one cysteine and two aspartic proteases), one unclassified desiccation-related protein, one cytoplasmic/mitochondrial alcohol dehydrogenase, one plasma membrane leucine rich repeat protein, three secretory peroxidases, one plasma membrane xylosidase, a secretory esterase/lipase, one secretory nuclease, and two amylases^13^. Of these, the amylase had not been previously described in the digestive fluids of other carnivorous plants and may suggest that non-defense related genes have also been coopted for carnivory.

The position of the Lentibulariaceae has yet to be resolved. The family has been placed sister to the Stilbaceae^1^, Schlegeliaceae^14^, Pedaliaceae and Acanthaceae^15^, and Acanthaceae^16–18^ using limited nuclear and plastid gene sets.

Here we present a chromosome-level assembly for *Pinguicula gigantea*, the second chromosome-level assembly for carnivorous plants. We also produced high quality assemblies for *Pinguicula agnata* and *Pinguicula moctezumae* primarily from long-read Oxford Nanopore Technologies (ONT) reads. We show evidence that *Pinguicula* has undergone an additional whole genome duplication beyond the duplication event for all core Lamiales. We also show that *Pinguicula gigantea* and the two long-read assemblies contain multiple putative tandem duplications for candidate digestive enzyme genes. Among these are tandemly duplicated cysteine proteases that are syntenic to two tandem duplications in *Utricularia gibba* with trap specific expression. Notably, these tandem cysteine protease genes appear to have independently duplicated in both *Pinguicula* and *Utricularia gibba*. Lastly, we place the Lentibulariaceae sister to the Acanthaceae, within the Lamiales.

## Results

### DNA Sequencing

Oxford Nanopore Technologies (ONT) sequencing of *Pinguicula gigantea* resulted in 8,977,265 reads with an N50 of 35,854 and a total of 55,499,618,859 bases. *Pinguicula agnata* had 12,613,739 reads with an N50 of 17,455 and a total of 39,854,418,704 bases. *Pinguicula grandiflora* had 4,683,883 reads with an N50 of 37,878 and a total of 28,445,068,544 bases. *Pinguicula moctezumae* had 12,121,023 reads with an N50 of 37,492 and a total of 52,762,913,837 bases. All ONT sequencing statistics are in Table S2.

Paired-end Illumina reads for the same samples were 251 bp in length. *Pinguicula agnata* had 38,806,442 paired-end reads, *Pinguicula gigantea* had 34,842,842, *Pinguicula grandiflora* had 40,266,500, and *Pinguicula moctezumae* had 37,551,390 (Table S3). Nanodrop and QUBIT measurements for extractions for RNA-seq reads can be found in Table S4. Paired-end Illumina reads for these four species were sequenced on the Illumina NextSeq 550 platform.

### Genome assembly

The primary assembly for *Pinguicula gigantea* had 884 contigs, 97.5% complete BUSCOs and 20.8% duplicated BUSCOs (Table S5). The haploid assembly generated by Purge Haplotigs has 405 contigs, 96.9% complete BUSCOs, and 15.1% duplicated BUSCOs (Table S5). For Hi-C scaffolding, 66,448,820 read pairs were generated for the Dovetail Hi-C library. A single break was made in the input assembly, and 401 joins were made. The assembly was scaffolded into 11 pseudomolecules with 10 additional small scaffolds (Figure 1). This Hi-C assembly had 96.8% complete BUSCOs and 14.3% duplicated BUSCOs (Table S5). The final assembly size was 608.7 Mbp, which is very close to the size estimated by flow cytometry, 598 Mpb^19^.

**Figure 1:**
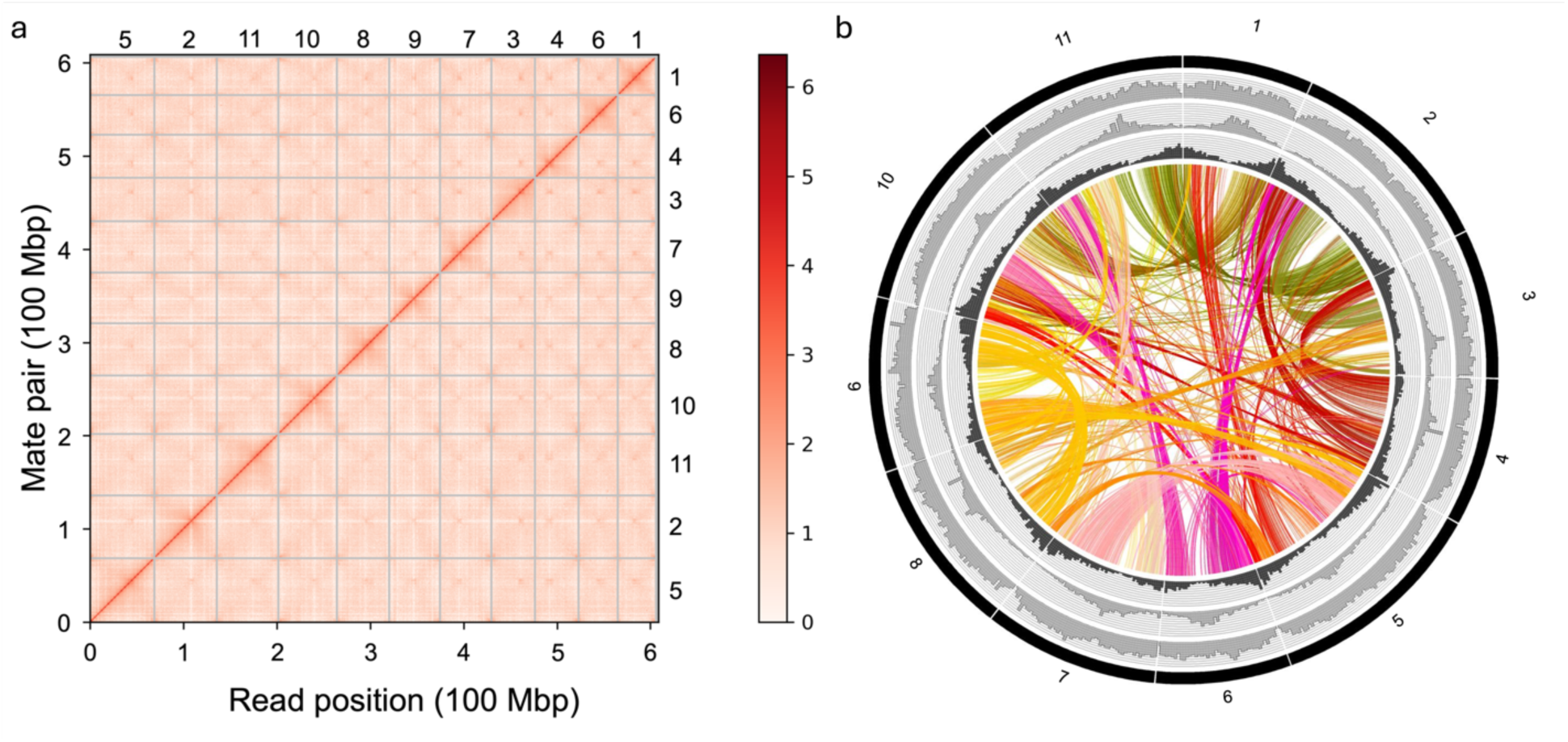
Chromosome-Scale Architecture of the *Pinguicula gigantea* Genome. (a) Darker red indicates a stronger signal. Read position on the Left and bottom. Chromosome numbers are on the right and top. (b) From the outer to the inner ring, the tracks represent: Chromosome number, LTR/Copia repeat density, LTR/Gypsy repeat density, and gene density. Putative centromeric regions are highlighted by peaks of increased transposable element (TE) density and decreased gene density. The plot also includes self-self syntenic connections, illustrating regions of genomic similarity.

The primary assembly for *Pinguicula grandiflora* had 1879 contigs, 95.1% complete BUSCOs, and 49.4% complete BUSCOs (Table S5). Each attempt at creating a haploid assembly reduced the BUSCO score too greatly, so the primary assembly was used as input for Hi-C scaffolding. For Hi-C scaffolding, 62,393,800 read pairs were generated for the Dovetail Hi-C library. Seven breaks were made in the input assembly, and 369 joins were made (Figure S1). The 1,879 contigs of the input assembly were scaffolded into 1,517 pieces. The final assembly size was 570.7 Mbp, which is larger than the size estimated by flow cytometry, 424 Mpb^19^. Due to quality issues (also mentioned below regarding syntenic gene ratios), the Hi-C assembly for *Pinguicula grandiflora* and its associated de novo gene annotation were only used for homology-based gene prediction, which was carried out using Gene Model Mapper (GeMoMa).

Two additional assemblies were generated from the ONT reads for *Pinguicula agnata* and *Pinguicula moctezumae*. Each of these assemblies had a high level of completeness based on BUSCO scores, but also a high number of duplicated BUSCOs (Table S6), which was likely due to uncollapsed heterozygosity as is common with long-read assemblies^20^. Using Purge Haplotigs, the number of duplicated BUSCOs decreased (Table S6): 14.2% for *Pinguicula agnata* and 15.6% for *Pinguicula moctezumae*, both very close to the percent of duplicated BUSCOs found for *Pinguicula gigantea* (Table S5). The final assembly sizes for *Pinguicula agnata* and *Pinguicula moctezumae* were 676.0 Mbp and 586.2 Mbp, which is marginally larger than the sizes estimated by flow cytometry, 651 Mbp and 572 Mbp, respectively^19^.

Both long-read assemblies were scaffolded using the *Pinguicula gigantea* reference assembly as a guide. The reference assembly was only used as a scaffolding guide and did not fill gaps with its own sequences between scaffolded contigs. *Pinguicula agnata* and *Pinguicula moctezumae* had 98.89% and 98.51% of their bp placed into scaffolds, respectively (Table S7).

### Genome annotation

The de novo gene annotation for the *Pinguicula gigantea* Hi-C assembly contained 35,940 genes, 92.3% complete BUSCOs, and 11.0% duplicated BUSCOs (Table S5). The de novo gene annotation for the *Pinguicula grandiflora* Hi-C assembly contained 33,294 genes, 88.8% complete BUSCOs, and 33.8% duplicated BUSCOs (Table S5). All other annotations generated with Gene Model Mapper (GeMoMa). Results for primary and purged assemblies used for Hi-C are found in Table S5. Results for the long-read assemblies are found in Table S6. Results for the scaffolded assemblies are found in Table S7. BUSCO scores for gene annotations correlated closely with the assembly BUSCO scores.

Repeat annotations for the short-read assemblies that were scaffolded with RagTag are found in Table S8. Notably, Ty1/Copia LTR retrotransposons made up the largest percentage of the genome for the *Pinguicula gigantea* reference assembly and the scaffolded long-read *Pinguicula agnata* and *Pinguicula moctezumae* assemblies.

The Hi-C assembly for *Pinguicula grandiflora*, which is in a separate clade from the other assemblies^21^, show a different pattern in repeat content (Table S8). *Pinguicula grandiflora* has the highest percentage of its genome consisting of Gypsy/DIRS1 LTR retrotransposons at 15.31% and unclassified repeats at 43.09%, two to three-fold more than the other assemblies.

### BUSCO trees

A BUSCO tree was generated for a group of thirty Lamiales genome assemblies and two Gentianales genome assemblies. Every BUSCO protein with four or more assemblies represented was used to generate a species tree. This resulted in 2,322 individual BUSCO protein trees that were used as an input for ASTRAL to generate a consensus tree (Figure 2a).

**Figure 2:**
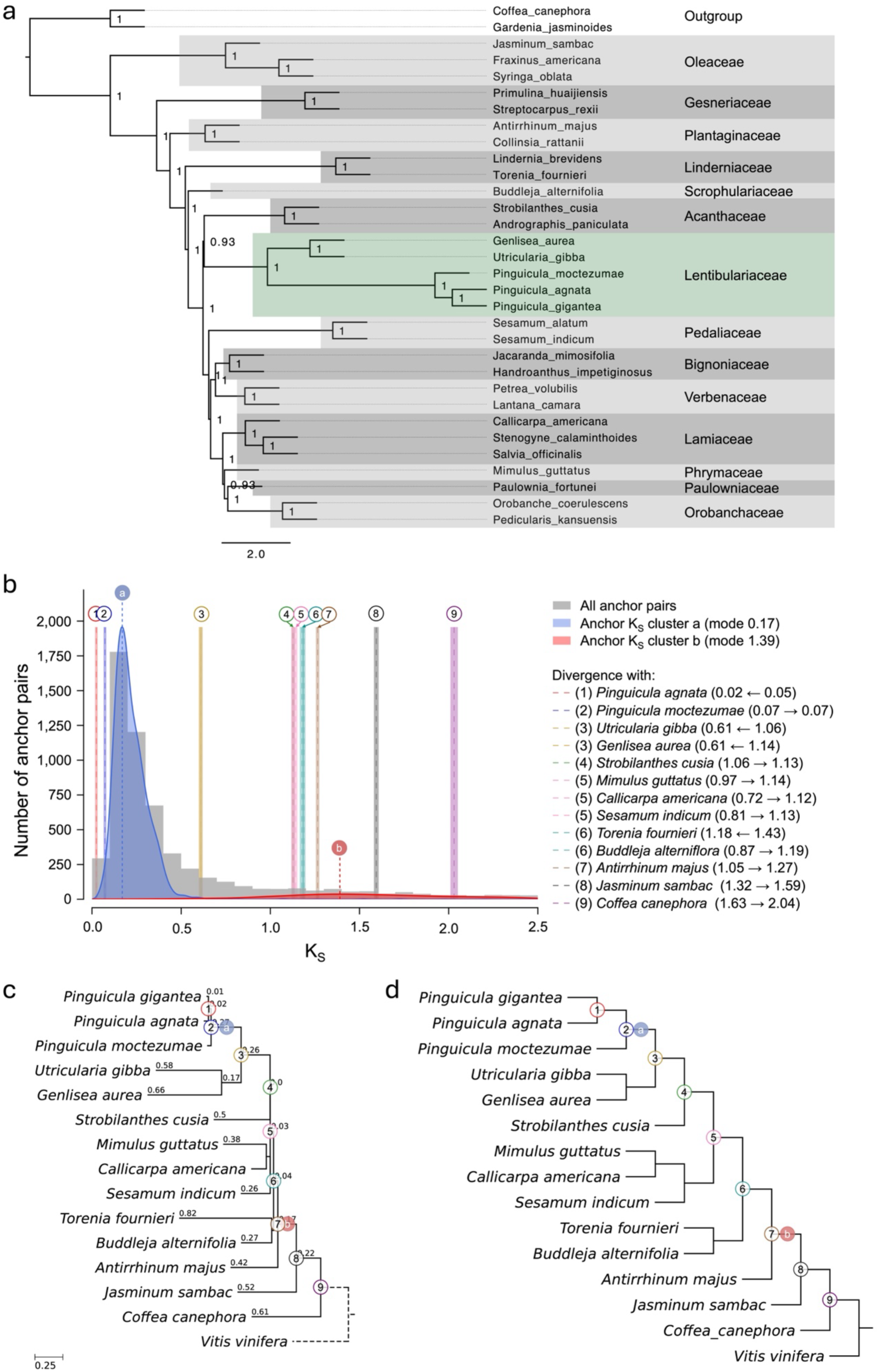
Phylogenomic placement and polyploid history of the Lentibulariaceae. (a) The Lentibulariaceae is highlighted in green. Other families of the Lamiales are highlighted in dark and light shades of grey. Two species from the Rubiaceae (Gentianales) were used as the outgroup. Tree generated from 2,322 BUSCO protein sequences. Branch support is local posterior probability. (b) Positioning of divergence times in the Lamiales relative to polyploidy events in *Pinguicula gigantea*. Syntenic (anchor) Ks parolog pair distribution for *Pinguicula gigantea* is shown in gray. Lognormal mixture model clustering of the median Ks values of the syntenic paralog pairs indicate the core Lamiales WGD in red, and a *Pinguicula*-specific WGD in blue. Vertical dashed lines labeled a and b denote the modes of these curves. Vertical lines labeled with 1–9 are rate-adjusted modal estimates of one-to-one ortholog Ks distributions between *Pinguicula gigantea* and other species, representing divergence times. (c) Phylogenetic tree branch lengths represent Ks distances estimated from ortholog Ks distributions. Dashed branches have lengths that cannot be computed by the software. Manually superimposed labels match with the Ks plot and represent *Pinguicula gigantea* polyploidy events and divergence with other lineages on the tree. (d) Same as (c), but with meaningless branch lengths to help see tree topology.

### Syntenic Mapping, Ks Plots, Fractionation Bias

The *Pinguicula gigantea* reference assembly has strong internal synteny between large blocks of paralogous genes, resulting from the most recent whole genome duplication (WGD) (Figure S2). Additionally, when the values for synonymous substitution rates (Ks) between each syntenic gene pair are plotted in a histogram, very few have low values, suggesting that this genome contains very little uncollapsed heterozygous sequences (Figure S2).

The *Pinguicula gigantea* reference assembly was also compared against *Vitis vinifera* as a quality check. Syntenic genes between *Pinguicula gigantea* and *Vitis vinifera* showed a 4:1 ratio (Figure S3c), suggesting two rounds of WGD in *Pinguicula gigantea*’s evolutionary history since it diverged from *Vitis vinifera*. For example, *Vitis vinifera* chromosome 1 matched with four large syntenic blocks in *Pinguicula gigantea* (Figure S3a). A similarly clear pattern is seen with *Vitis vinifera* chromosome 16 (Figure S3a). This can also be observed in the fractionation bias plots. *Pinguicula gigantea* has two sets of two chromosome sequences with similar patterns of gene retention compared to a single chromosome of *Vitis vinifera* (Figure S3b,d).

*Pinguicula gigantea*’s genome was also compared against *Mimulus guttatus* in the same way it was compared to *Vitis vinifera*. Syntenic genes between *Pinguicula gigantea* and *Mimulus guttatus* showed a 4:2 ratio (Figure S4c), suggesting one additional WGD in *Pinguicula gigantea*’s evolutionary history since it diverged from *Mimulus guttatus*. For example, a segment of *Mimulus guttatus* chromosome 11 matched with four large syntenic blocks in *Pinguicula gigantea* (Figure S4a). At the same time, each of those four syntenic blocks in *Pinguicula gigantea* is represented twice in *Mimulus guttatus*’ genome (Figure S4a). A similarly clear pattern is seen with *Mimulus guttatus* chromosome 3 (Figure S4a). This can also be observed in the fractionation bias plots. *Pinguicula gigantea* has a set of two chromosome sequences with similar patterns of gene retention compared to a single chromosome of *Mimulus guttatus* (Figure S4b,d).

For the *Pinguicula grandiflora* Hi-C assembly, when compared against *Pinguicula gigantea*, there was a 3:2 syntenic gene ratio (Figure S5). The expected syntenic gene ratio was 4:2 since *Pinguicula grandiflora* (4n=32^22^) is believed to have doubled one more time than *Pinguicula gigantea* (2n=22^22^). Due to uncertainty about this Hi-C assembly, this version of the *Pinguicula grandiflora* assembly was not used beyond an input reference for homology-based gene prediction was carried out using GeMoMa.

For the two primary long-read assemblies, when compared against *Pinguicula gigantea* using a syntenic dot plot map, *Pinguicula agnata* had a 2:4 syntenic gene ratio (Figure S6a,b) and *Pinguicula moctezumae* had a 2:3 syntenic gene ratio (Figure S7a,b). Additionally, when these genomes were analyzed for internal synteny between paralogous gene pairs, many of these syntenic gene pairs had Ks values that were much lower than observed in the *Pinguicula gigantea* reference assembly (Figure S8a,b & S9a,b). *Pinguicula agnata* and *Pinguicula moctezumae* are thought to only share the same polyploid events as *Pinguicula gigantea*, so these low Ks values are likely uncollapsed heterozygous sequences. After using PurgeHaplotigs, many of these low Ks syntenic gene pairs were removed (Figure S8c,d & S9c,d). Additionally, when these purged assemblies were compared against *Pinguicula gigantea* using a syntenic dot plot map, both *Pinguicula agnata* (Figure S6c,d) and *Pinguicula moctezumae* (Figure S7c,d) had a 2:2 syntenic gene ratio against *Pinguicula gigantea*.

The syntenic depth of the *Pinguicula gigantea* reference assembly was also compared against *Utricularia gibba*. By altering the maximum number of nearby genes searched to find at least five gene pairs, the syntenic depth between *Pinguicula gigantea* and *Utricularia gibba* was 2:3 (searching twenty nearby genes; Figure S10a) and 2:6 (searching fifty nearby genes; Figure S10b). This indicates that *Utricularia gibba* has undergone more duplication events^8,23^ than *Pinguicula gigantea* since diverging from one another.

### Tandem digestive enzyme gene analysis

The genome assembly of *Pinguicula gigantea* contains tandemly duplicated cysteine protease genes (Figure 3). This tandem duplication is located on chromosome 8, but not within the paralogous syntenic block on chromosome 5, suggesting the tandem duplications occurred after the most recent whole genome duplication in *Pinguicula gigantea*’s evolutionary history. These syntenic blocks in *Pinguicula gigantea* are also syntenic to both of *Utricularia gibba*’s cysteine protease tandem duplications on sister unitigs 699 and 85(+unitig 27)^8^ (Figure S11). The genome assemblies of *Pinguicula agnata* and *Pinguicula moctezumae* also each contain two regions that are syntenic with the *Pinguicula gigantea* tandem duplication. In each case, one syntenic block contains a tandem duplication for the cysteine protease genes and the other lacks a tandem duplication (Figure S12).

**Figure 3:**
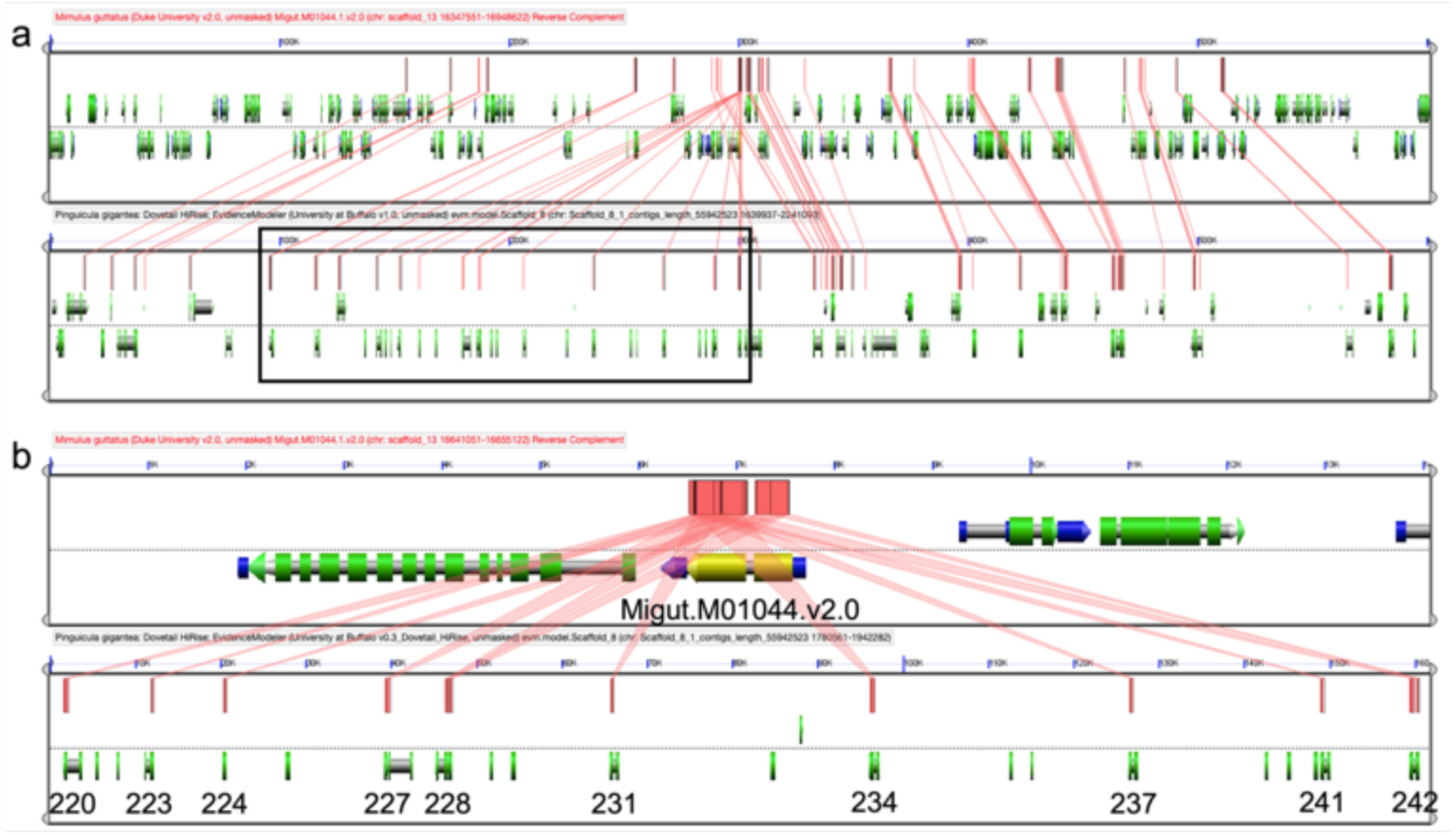
Cysteine protease tandem array in *Pinguicula gigantea* and homologous region in *Mimulus guttatus*. (a) Syntenic blocks between *Mimulus guttatus* scaffold 13 and *Pinguicula gigantea* chromosome 8. The tandem array is marked with a black rectangle. (b) Zoom in of the tandem array. Gene models are labeled with the full gene name in *Mimulus* above and just with the gene number on chromosome 8 in *Pinguicula* below. CDS are represented by green rectangles. Colored lines connect homologous gene pairs.

In all, eighteen tandemly duplicated putative cysteine protease genes on chromosome 8 were identified by CoGe Blast using BAW35427.1 as the search query (Table S9). Using InterPro^24^ to compare protein domains between the *Pinguicula gigantea* cysteine proteases and those from *Arabidopsis thaliana*, *Drosera adelae*, and *Utricularia gibba*, there were differences in the presence and absence of protein families, domains, homologous superfamilies, active sites, and conserved site (Table S10). Notably, only three *Pinguicula gigantea* proteins contained IPR000169, the cysteine active site which is part of a catalytic triad along with IPR025660 and IPR025661 (https://www.ebi.ac.uk/interpro/entry/InterPro/IPR000169/)^24^. The three *Pinguicula gigantea* proteins with this active domain were missing IPR025661 or both IPR025660 and IPR025661.

The cysteine protease coding sequences (CDS; Figure S13a) and proteins (Figure S13b) were also used to generate trees. Notably, both *Utricularia gibba* and *Pinguicula gigantea* have many sequences that cluster together by species. In the CDS tree, most of the *Pinguicula* and *Utricularia* tandem genes are sister to each other. Some genes from the tandem array group separately from the others. Pgig_chr8.220 and Pgig_chr8.242 separately group with the two genes from the paralogous syntenic block without the tandem array, Pgig_chr5.3731 and Pgig_chr5.3731, respectively. The protein tree further separates the *Pinguicula* tandem cluster from the *Utricularia* tandem clusters, besides Pgig_chr8.242 and Pgig_chr5.3730 grouping closer to the *Utricularia gibba* tandem genes than of the *Pinguicula gigantea* tandem genes. Similarly to the CDS tree, Pgig_chr5.3731 groups separately from the other tandem genes with Pgig_chr8.220. CDS and protein trees were generated for all digestive enzymes identified in the digestive fluid of *Pinguicula x Tina*. The focus was on the genes with three or more tandem duplicates in the *Pinguicula gigantea* reference assembly, but none had as expansive tandem duplication as the cysteine proteases.

Additional putative tandem gene duplications were identified for other candidate digestive enzyme genes in *Pinguicula gigantea*. These were for aspartic protease (Figure S14), peroxidase 10-like (Figure S15), putative peroxidase (Figure S16), and predicted esterase/lipase (Figure S17, Table S9). Tandems were not identified for putative beta-xylosidase (Figure S18), desiccation-related protein (Figure S19), or alpha-amylase genes (Figure S20).

## Discussion

Here, we present a high quality, reference genome for *Pinguicula gigantea*. This is the second chromosome-level genome assembly for a carnivorous plant^12^ and will be instrumental in comparative genome analysis with other carnivorous plants, especially with other carnivorous genera within the Lentibulariaceae, *Utricularia* and *Genlisea*. Using this genome assembly, we confirmed the polyploidy events in the evolutionary history of *Pinguicula*. Comparing syntenic blocks between *Pinguicula gigantea* and *Vitis vinifera* clearly shows two whole genome duplication (WGD) events in the evolutionary history of *Pinguicula gigantea* since splitting from *Vitis vinifera* (Figure S3). The first WGD is shared among the core Lamiales ^15^ and the second is plausibly limited to a subset of *Pinguicula* species (figure 2b-d). The second WGD can also be seen specifically when comparing *Pinguicula gigantea* against *Mimulus guttatus* (Figure S4), which only has experienced a single WGD in its evolutionary history since diverging from *Vitis vinifera*^15,23^. These assemblies show a 4:2 pattern between syntenic genes, indicating that after these two species diverged, the ancestor of *Pinguicula gigantea* underwent an additional WGD. This genome also aided in determining the polyploidy events in *Utricularia gibba*. Using the original 81.87 Mbp assembly, *Utricularia gibba* was thought to have undergone at least three WGD events since the gamma hexaploidy event about 120 MYA^23^. Later, with an improved 102 Mbp assembly, *Utricularia gibba* was now proposed to have at least two and as many as three WGD events since the gamma event^8^. The One Thousand Plant Transcriptomes Initiative concluded that *Utricularia* and *Pinguicula* only shared the core Lamiales WGD and had their own separate WGD events, PICA𝛼 and UTRI𝛼^17^, inferred by both Ks plots and synonymous ortholog divergence analyses. When the *Pinguicula gigantea* reference assembly was compared with the *Utricularia gibba*, syntenic blocks between the two assemblies show between a 2:3 and 2:6 pattern (Figure S10), suggesting that *Utricularia gibba* has undergone more WGD events^8,23^ than *Pinguicula gigantea* has since the shared core-Lamiales WGD event.

We also produced two highly contiguous and highly complete de novo long-read assemblies (Table S6). *Pinguicula agnata* and *Pinguicula moctezumae* both also had a 2:2 syntenic gene ratio against the *Pinguicula gigantea* reference assembly, which was as expected based on their shared ploidy level, 2n=22^22^ (Figure S6c,d & S8c,d).

We found that the Lentibulariaceae, the family of *Pinguicula*, *Utricularia*, and *Genlisea*, was sister to the Acanthaceae (Figure 2a). This supports its position in previously published phylogenies^16–18^. Due to a lack of published nuclear genomes from the Stilbaceae and Schlegeliaceae, the relationships between either family and the Lentibulariaceae could not be examined. Compared with the *Pinguicula* clade’s relatively short branch lengths, long branch lengths were found for the *Utricularia* and *Genlisea* clade, which had been reported previously^25^. These differences are likely due to higher mutation rates in *Utricularia* and *Genlisea* compared with *Pinguicula*^26,27^

The *Pinguicula gigantea* genome contains a putative tandem duplication of cysteine protease genes (Figure 3). We currently lack transcriptional evidence for enhanced transcription of cysteine proteases in trap tissues like what was found in *Utricularia gibba*^8^. This is primarily due to the body plan of *Pinguicula*, which lacks non-trap photosynthetic tissues. When we compare the InterPro^24^ matches between the cysteine protease proteins in *Utricularia gibba*^8^, the *Pinguicula gigantea* reference in this study, *Drosera Adelae* (BAW35427.1)^28^, and *Arabidopsis thaliana* (AT5G45890.1)^29^, there are some noteworthy comparisons (Table S10). The sequences in *A. thaliana*, *D. adelae*, *Utricularia gibba*, and *Pinguicula gigantea* contained the same InterPro family (IPR013128), domains (IPR013201, IPR000668, IPR039417), and homologous superfamily (IPR038765). Additionally, *A. thaliana* and most of the proteins from *Utricularia gibba* contained three active sites identified by InterPro (IPR025660, IPR025661, IPR000169). The sequence from *D. adelae* only contained two of the three active sites (IPR025661, IPR000169). No protein sequence from *Pinguicula gigantea* contained all three active sites and only two with all the family, domain, and homologous superfamily IDs mentioned above contained active site IPR000169 (Pgig_chr8.220, Pgig_chr8.228). Having all three active sites is not necessary for transcription. Between the tandem duplicates on unitig 699 and 85, Lan et al.^8^ found four genes with high levels (unitig_85.g27286, unitig_85.g27287, unitig_699.g19346, unitig_699.g19347) and five genes with more moderate levels of trap-specific transcription (unitig_85.g27288, unitig_85.g27291, unitig_85.g27292, unitig_699.g19345, unitig_699.g19348). Two of the four highly expressed proteins lack one active site (unitig_85.g27286.t1, unitig_699.g19347.t1) and two of the five proteins with more moderate levels of expression lack one active site (unitig_699.g19345.t1, unitig_699.g19348.t1).

Based on the results from the cysteine protease CDS and protein trees (Figure S13), *Pinguicula* and *Utricularia* evolved tandem arrays of cysteine protease genes independently. This result comes from the fact that all the *Utricularia gibba* tandem sequences cluster separately from the *Pinguicula* sequences, suggesting that they evolved separately instead of from a common ancestor. While it’s possible that these genes only artificially appear to be more closely related to near-by paralogous sequences through gene conversion^30^, we did not test for possibility. Assuming that the trees are true, we propose the following order of events: *Pinguicula* and *Utricularia* diverged from each other, and each lineage experienced a WGD (Figure 2b-d).

Following this WGD, *Pinguicula* independently experienced a tandem duplication of cysteine proteases in one region of its genome. After diverging from *Pinguicula*, *Utricularia gibba* likely experienced one WGD, evolved the tandem array of cysteine proteases, and underwent an additional WGD, doubling its number of tandem arrays of cysteine protease genes relative to *Pinguicula gigantea* (Figure S11). *Pinguicula agnata*, *Pinguicula gigantea*, and *Pinguicula moctezumae* did not experience additional polyploidy events, meaning that these species all have one syntenic region with the tandem duplication and one without (Figure S11 & S12). Notably, the tandem array of cysteine protease genes in *Pinguicula gigantea* continued to experience more tandem duplication than the others (Figure S12).

Because the cysteine protease gene cluster in *Utricularia gibba* is located within regions of LARDs, a type of retrotransposon^8^, we examined the repeat annotation that were in and around Pgig_chr8.218 through Pgig_chr8.242 (chromosome 8, positions 1,756,076-1,942,842). We determined that unknown elements made up the largest percentage of the number of elements at 63.9%, LTR/Copia were 10.4%, simple repeats were 10.1%, LINE/L1 were 7.6%, LTR/Unknown were 3.7% and low complexity were 2.1% (Table S11). LTR/Gypsy made up 0.9% of the elements, tRNA made up 0.6%, rRNA made up 0.3%, and DNA/Zisupton made up 0.3% (Table S11). As noted in Lan et al.^8^, these repeat elements may have facilitated the tandem duplication events un *Utricularia gibba* and *Pinguicula*. When compared against repeat elements within its tandemly duplicated regions of cysteine proteases on scaffolds 8, 85, 578, and 699 of *Utricularia gibba*, 63.7% were unknown, followed by 22.6% simple repeats, 4.2% low complexity, 3.0% LTR/Copia, 3.0% DNA/hAT-Ac, 1.2% LTR/Gyps, 1.2% DNA/PIF-Harbinger, 0.6% tRNA, and 0.6% DNA (Table S12). Careful, manual annotation of these tandemly duplicated regions, along with transcriptome data may be necessary to fully confirm these regions in the future.

*Pinguicula gigantea* contained tandem duplications for many of the other putative digestive enzymes, although these were smaller than the cysteine protease duplication. Notably, there were no tandem duplications for the putative beta-xylosidase (Figure S18), desiccation-related protein (Figure S19), or alpha-amylase genes (Figure S20). If any of these enzymes were important for the evolution of the carnivorous syndrome in *Pinguicula gigantea*, a mechanism other than tandem gene duplication would have been requisite.

## Conclusions

Here, we present a chromosome-level assembly for the carnivorous plant *Pinguicula gigantea*. Using this genome in conjunction with three additional long-read assemblies, we uncovered the polyploid history of *Pinguicula* and have a better understanding of its close relative, *Utricularia gibba*, as well. We also place the Lentibulariaceae sister to Acanthaceae using the most sequence data to date. Finally, we discovered a large tandem array of cysteine protease genes that are homologous with a tandem duplication of cysteine protease genes with trap specific expression in *Utricularia gibba*. We suggest that these two tandem arrays evolved independently, with different collections of surrounding repeat elements, potentially facilitating the duplication process.

## Methods

### Material collection and DNA Sequencing

Oxford Nanopore Technologies (ONT) reads for *Pinguicula agnata*, *Pinguicula gigantea*, *Pinguicula grandiflora*, and *Pinguicula moctezumae* were provided by Todd P. Michael of the J. Craig Venter Institute, California, USA. Illumina paired end reads generated on the Illumina NextSeq 550 platform for the same individuals at The Pennsylvania State University. Corrupted ONT reads for *Pinguicula agnata* were removed using fastqwiper version 1.0.5-alpha (https://github.com/mazzalab/fastqwiper). ONT read quality was assessed using NanoPlot version 1.42.0^31^. Illumina read quality was assessed with fastqc version 0.11.9 ^32^. RNA-seq reads were generated using NextSeq High Output 150 x 150 paired-end sequencing.

### Genome assembly

De novo Genome assemblies for *Pinguicula agnata*, *Pinguicula gigantea*, *Pinguicula grandiflora*, and *Pinguicula moctezumae* utilized both Oxford Nanopore Technologies (ONT) long read data and Illumina short read data. The genome assembly for *Pinguicula gigantea* was carried out using the Minimap2 version 2.16 and Miniasm version 0.3-r179^33,34^ de novo assembly pipeline. Racon version 1.3.3^35^ was run three times to produce a consensus assembly and Pilon version 1.23^36^ was run three times for error correction using Illumina short read data. Genomes were indexed using SAMtools version 0.1.19^37^ and BWA version 0.7.17-r1188^38^, ONT long reads were mapped against the genome using Minimap2 version 2.16, Illumina short reads were mapped against the genome using bwa version 0.7.17-r1188, and the resulting BAM files was sorted using SAMtools version 0.1.19. Purge Haplotigs version 1.1.1^39^ was used to generate a haploid genome assembly.

For *Pinguicula gigantea*, a Dovetail Hi-C library was prepared in a similar manner as described previously^40^. Briefly, for each library, chromatin was fixed in place with formaldehyde in the nucleus and then extracted fixed chromatin was digested with DpnII, the 5’ overhangs filled in with biotinylated nucleotides, and then free blunt ends were ligated. After ligation, crosslinks were reversed, and the DNA purified from protein. Purified DNA was treated to remove biotin that was not internal to ligated fragments. The DNA was then sheared to ∼350 bp mean fragment size and sequencing libraries were generated using NEBNext Ultra enzymes and Illumina-compatible adapters. Biotin-containing fragments were isolated using streptavidin beads before PCR enrichment of each library. The libraries were sequenced on an Illumina HiSeq X to a target depth of 30x coverage.

The input de novo assembly and Dovetail Hi-C library reads were used as input data for HiRise, a software pipeline designed specifically for using proximity ligation data to scaffold genome assemblies^41^. Dovetail Hi-C library sequences were aligned to the draft input assembly using a modified SNAP read mapper^42,43^. The separations of Dovetail Hi-C read pairs mapped within draft scaffolds were analyzed by HiRise to produce a likelihood model for genomic distance between read pairs, and the model was used to identify and break putative misjoins, to score prospective joins, and make joins above a threshold.

*Pinguicula grandiflora* was also assembled in the same manner as *Pinguicula gigantea*, except with ONT reads shorter than 10k bp filtered out using NanoFilt version 2.5.0^31^ and without a Purge Haplotigs step. The draft assembly was then sent to Dovetail for Hi-C scaffolding. For the Dovetail Omni-C library, chromatin was fixed in place with formaldehyde in the nucleus and then extracted. Fixed chromatin was digested with DNAse I, chromatin ends were repaired and ligated to a biotinylated bridge adapter followed by proximity ligation of adapter containing ends. After proximity ligation, crosslinks were reversed and the DNA purified. Purified DNA was treated to remove biotin that was not internal to ligated fragments. Sequencing libraries were generated using NEBNext Ultra enzymes and Illumina-compatible adapters. Biotin-containing fragments were isolated using streptavidin beads before PCR enrichment of each library. The library was sequenced on an Illumina HiSeqX platform to produce ∼ 30x sequence coverage. Then HiRise used (See read-pair above) MQ>50 reads for scaffolding.

The input de novo assembly and Dovetail OmniC library reads were used as input data for HiRise, a software pipeline designed specifically for using proximity ligation data to scaffold genome assemblies^41^. Dovetail OmniC library sequences were aligned to the draft input assembly using bwa (https://github.com/lh3/bwa). The separations of Dovetail OmniC read pairs mapped within draft scaffolds were analyzed by HiRise to produce a likelihood model for genomic distance between read pairs, and the model was used to identify and break putative misjoins, to score prospective joins, and make joins above a threshold. The final Hi-C genome for *Pinguicula grandiflora* was annotated using the same de novo method as *Pinguicula gigantea* (described below in the annotation section). This *Pinguicula grandiflora* genome assembly was not included in structural analyses due to quality issues, but it was used as a reference for GeMoMa annotations described below.

The genome assemblies for *Pinguicula agnata* and *Pinguicula moctezumae* were carried out using the Minimap2 version 2.24 and Miniasm version 0.3-r179^33,34^ de novo assembly pipeline. Racon version 1.5.0^35^ was run five times to produce a consensus assembly and Pilon version 1.23^36^ was run three times for error correction using Illumina short read data. Genome assemblies were indexed using SAMtools version 1.16.1 and BWA version 0.7.17-r1188, ONT long reads were mapped against the genome using Minimap2 version 2.24, Illumina short reads were mapped against the genome using bwa version 0.7.17-r1188, and the resulting BAM files was sorted using SAMtools version 1.16.1. Repetitive element annotations (described below in the annotation section) were converted into a BED-format file for input in Purge Haplotigs so those sequences would be ignored during Purge Haplotigs. Purge Haplotigs version 1.1.2 ^39^ was used to generate haploid genome assemblies.

The two purged MiniMap/Miniasm assemblies were scaffolded into pseudo-chromosome-level assemblies using RagTag version 2.1.0^44^ and our chromosomal *Pinguicula gigantea* reference assembly as a reference. Assembly statistics were generated using Quality Assessment Tool for Genome Assemblies (QUAST) version 5.0.2^45^, and assembly completeness was measured using Benchmarking Universal Single-Copy Orthologs (BUSCO) version 5.5.0 embryophyta_odb10^46^.

### Genome annotation

Genome annotations for *Pinguicula gigantea* and *Pinguicula grandiflora* were generated by combining transcriptome assembly and gene prediction data using EVidenceModeler version 1.1.1^47,48^. RNA-seq reads were trimmed using trimmomatic-0.39^49^. Repeat content was characterized and masked using RepeatModeler version 2.0.1^50^ and RepeatMasker version 4.1.0^51^ run with the rmblast search engine version 2.10.0 (https://www.repeatmasker.org/rmblast/). In addition to the default RepeatScout (version 1.0.6)^52^ /RECON (1.0.8)^53^ pipeline, the LTR (Long Terminal Repeat) structural discovery pipeline was run as well for RepeatModeler using LtrHarvest (GenomeTools version 1.5.9)^54^ and LTR_retriever version 2.9.0^55^. Four independent transcriptome assemblies were carried out using Trinity version 2.12.0^56–58^, HISAT2 version 2.2.1^58^ and StringTie version 2.1.5^59^, STAR version 2.7.9^60^ and StringTie, and TransAbyss version 2.0.1^61^.

The four transcriptomes were combined and filtered using EvidentialGene (July 30, 2021 download)^62^. Program to Assemble Spliced Alignments (PASA) version 2.5.1^63^ was used to generate a gff3 annotation file from the merged transcriptome using the parallelized BLAST-like alignment tool (pblat) version 2.5^64^, Genomic Mapping and Alignment Program (GMAP) (August 25, 2021 download)^65^, and Minimap2 version 2.20 aligners. Homology-based gene prediction was carried out using Gene Model Mapper (GeMoMa) version 1.7.1^66,67^ using default settings and five different species as references: *Utricularia gibba*^8^, *Mimulus guttatus*^68^, *Tectona grandis*^69^, *Sesamum indicum*^70^, and *Callicarpa americana*^71^. Furthermore, gene models were independently predicted against the masked genome assembly using GeneMark-ES version 4.68 ^72^ and BRAKER version 2.1.6^37,72–78^ (mapping done with STAR version 2.7.9 and sorted using SAMtools version 1.11^37^). The PASA, BRAKER, GeneMark, and GeMoMa gff3 annotation files were combined and filtered using EVidenceModeler with segment size set to 100,000 bp and overlap size set to 10,000 bp. Annotation completeness was measured by running BUSCO version 5.5.0 embryophyta_odb10.

All other genome annotations were carried out using Gene Model Mapper (GeMoMa) version 1.9 ^66,67^, a homology-based gene prediction software. Each GeMoMa annotation used the Dovetail *Pinguicula gigantea* and *Pinguicula grandiflora* assemblies with their de novo annotations as references. They also used *Utricularia gibba*^8^, *Genlisea aurea*^7^, *Mimulus guttatus*^68^, and *Arabidopsis thaliana*^29^ as references. Annotation completeness was measured by running BUSCO version 5.5.0 embryophyta_odb10 on GeMoMa’s predicted proteins file, followed by GeMoMa’s BUSCOrecomputer tool so BUSCO statistics were computed for genes instead of transcripts. Repeat content was characterized and masked using RepeatModeler version 2.0.1^50^ and RepeatMasker version 4.1.0^51^ run with the rmblast search engine version 2.10.0 (https://www.repeatmasker.org/rmblast/). In addition to the default RepeatScout (version 1.0.6)^52^ /RECON (1.0.8)^53^ pipeline, the LTR (Long Terminal Repeat) structural discovery pipeline was run as well for RepeatModeler using LtrHarvest (GenomeTools version 1.5.9)^54^ and LTR_retriever version 2.9.0^55^.

### BUSCO trees

BUSCO version 5.4.4 ^46^ was used for the construction of phylogenetic trees. BUSCOs were extracted from a select group of genome assemblies using the eudicots_odb10 lineage. Single-copy BUSCO protein sequences from each genome assembly were extracted and grouped by BUSCO using an in-house script. Each protein FASTA file was aligned with MAFFT version 7.490^79^ using default options and trimAl version 1.4.1^80^ using the automated1 option. IQ-TREE version 1.6.12^81^ was run on each individual BUSCO alignment that had 4 or more species included. IQ-TREE used 1000 SH-like approximate likelihood ratio test (SH-aLRT) replicates, 1000 Ultrafast bootstrap replicates^82^, and the best model was chosen for each tree using ModelFinder^83^. The resulting tree files were used as input for ASTRAL version 5.7.8^84^ with default options.

### Syntenic Mapping, Ks Plots, Fractionation Bias

Select genome assemblies and annotations were uploaded to the Comparative Genomics (CoGe) online platform^85^. Each assembly was input into SynMap2 ^86^ and run against itself to generate syntenic dot plot maps for paralogous genes and accompanying histograms for synonymous substitution rates (Ks) between syntenic gene pairs. This data was also downloaded for *Coffea canephora* (CoGe id), *Catharanthus roseus* (CoGe id65259), *Gelsemium elegans* (CoGe id64491), *Gelsemium sempervirens* (CoGe id53941), *Gardenia jasminoides* (CoGe id66926), *Ophiorrhiza pumila* (CoGe id63710), *Prunus persica* (CoGe id36122), *Theobroma cacao* (CoGe id64859), and *Vitis vinifera* (CoGe id19990). SynMap was also used to compare syntenic orthologous gene pairs between two species. In each case, SynMap’s Syntenic Path Assembly option ^87^ was utilized to help visualize and count the ratio of syntenic blocks between each pair of species. Assemblies were compared against *Vitis vinifera* (CoGe id19990)^88^, which has no additional polyploid events since the gamma hexaploidy event at the base of all core eudicots^88–90^, and *Mimulus guttatus* (CoGe id22665)^68^, which has had one whole genome duplication (WGD) shared with the other members of the core Lamiales since the gamma hexaploidy event^15^. Duplicate subgenomes were assessed to determine relative ploidy level using SynMap’s fractionation bias (FractBias) tool ^91,92^.

Select genome assemblies and annotations were uploaded to the Comparative Genomics (CoGe) online platform^85^. Each assembly was input into SynMap2^86^ and run against itself to generate syntenic dot plot maps for paralogous genes and accompanying histograms for synonymous substitution rates (Ks) between syntenic gene pairs. The Ks reports were downloaded for each run. Density plots (smoothed curves and ridge plots) of Ks values for syntenic paralogs were generated in R^93^ using the tidyverse^94^, ggplot2^95^, RColorBrewer^96^, ggridges^97^, ggpmisc^98^, summarizedpackages^99^, and data.table (DOI: 10.32614/CRAN.package.data.table).

Ksrates version 1.1.3 was used to position species splits relative to polyploidy events. Coding sequence (CDS) fasta files were extracted using AGAT version 0.9.2. Paralogous Ks peaks were generated using the genome assembly for *Pinguicula gigantea*. Orthologous Ks peaks were generated using combinations of species pairs for each species pair. Ks value of the *gamma* hexaploidy event (Figure 2b; green curve c) was justified by comparing the Ks peaks from multiple other species that have not undergone additional WGD since the *gamma* hexaploidy event (Figure S21).

### Tandem digestive enzyme gene analysis

Each of *Pinguicula x Tina*’s digestive enzyme protein sequences^13^ were input into CoGeBLAST^100^ to search for tandem duplicates in the *Pinguicula gigantea* reference assembly. All results were imported into a Fasta View and select results were imported into the Genome Evolution (GEvo) Analysis tool to compare syntenic blocks within and between genome assemblies.

MCScan^101^ was utilized to generate syntenic dot plots, syntenic depth, and macrosynteny karyotype plots between assemblies. Input annotation files were converted to gff3 format using AGAT version 0.9.2^102^. AGAT was also used to extract CDS fasta files using each species’ assembly and reformatted annotation. When detecting synteny between two species, default options were used to generate figures.

Digestive enzymes for gene trees were chosen using CDS and protein sequences identified by mass spectrometry in *Pinguicula x Tina* digestive fluid 24 hours after feeding on fruit flies^13^. NCBI BLAST+ tblastn and blastp version 2.12.0 ^103^ were used to search each CDS and protein database, respectively, against the enzyme protein sequence query. The following sequences were used as search queries: alpha-amylase (EPS60632.1 and KZV28895.1); aspartic protease (BAD07475.1, GAV80475.1); cysteine protease (AT5G45890.1, BAW35427.1); desiccation-related protein (BAW35440.1); peroxidase 10-like (KZV23101.1); predicted esterase lipase (XP004232991.1); putative beta-xylosidase (AAX92967.1); putative peroxidase (BAM28609.1). The resulting sequences were aligned with MAFFT version 7.490 ^79^ using default options and trimAl version 1.4.1 ^80^ using the automated1 option. The aligned and trimmed fasta file was used as input for IQ-TREE version 1.6.12 ^81^. IQ-TREE used 1000 SH-like approximate likelihood ratio test (SH-aLRT) replicates, and 1000 Ultrafast bootstrap replicates ^82^. CDS and protein trees incorporated sequences from the *Pinguicula gigantea* reference, the two long-read *Pinguicula* assemblies (*Pinguicula agnata* and *Pinguicula moctezumae*), and *Arabidopsis thaliana*^29^, *Callicarpa americana*^71^, *Coffea canephora*^104^, *Gardenia jasminoides*^105^, *Mimulus guttatus*^68^, *Sesamum indicum*, *Tectona grandis*^69^, *Utricularia gibba*^8^, and *Vitis vinifera*^88^.

We also used InterPro^24^ to examine the InterPro domains matches for each cysteine protease protein in the tandem cluster in *Utricularia gibba*^8^ and the *Pinguicula gigantea* reference generated in this study. We also included sequences from *Drosera adelae* (BAW35427.1)^28^ and *Arabidopsis thaliana* (AT5G45890.1)^29^.

### CIRCOS genome view

We first used the output from RepeatModeler^50^ to identify and isolate the tracks needed for the Gypsy and Copia density tracks (Figure 1b). Additionally, the *Pinguicula gigantea* genome annotation was filtered to isolate gene tracks. These tracks were then parsed to create histograms representing the density of Gypsy and Copia elements, as well as gene density, across the genome. The scaffold lengths were used to normalize the feature densities per Mbp. The data was then sorted to ensure accurate representation of scaffold order.

Self-self synteny lines were created by reformatting the Ks data output of a self-self synteny analysis produced by CoGe’s SynMap2^86^ tool. These lines illustrate regions of genomic similarity and were integrated into the visualization to provide a comprehensive view of the genomic architecture. Finally, the Circos software^106^ was used to combine these density tracks and synteny lines into a detailed Circos plot.

## Acknowledgments

VAA acknowledges funding from the United States National Science Foundation award 2030871 (“Genomic Architecture Underlying Biochemical Diversity and Molecular Convergence within the Carnivorous Plants”), through which Tanya Renner (Penn State University) was also funded and is gratefully acknowledged, including for production of RNA-seq reads used in this research.

## Authors’ Contributions

VAA conceived and led the project with important input from SJF; MMR, LH-E and EI-L provided plant tissue and/or other research materials; CT, TL and VAA prepared DNA; TPM sequenced DNA and produced primary genome assemblies; SJF, JK and SR annotated the genomes; SJF performed the majority of genome analyses (including reference-based genome scaffolding and Ksrates, CoGe, and enzyme phylogenetic analyses); RWJ, VAA, and SJF provided essential information on *Pinguicula* and relatives, JK generated the Circos plot and BUSCO species tree. SJF wrote the manuscript, with input from VAA and JK.

## Data availability

Original data obtained for this project is available at https://doi.org/10.5061/dryad.b8gtht7p2. DNA and RNA-seq reads are currently available on request, but they will be uploaded and released on the NCBI Short Read Archive when this research is formally published in a scientific journal.

## Notes

### Competing Interest Statement

The authors have declared no competing interest.

## References

1 Fleck, S. J. & Jobson, R. W. Molecular Phylogenomics Reveals the Deep Evolutionary History of Carnivory across Land Plants. Plants 12, 3356 (2023).

2 Fleischmann, A., Schlauer, J., Smith, S. & Givnish, T. in Carnivorous Plants: Physiology, ecology, and evolution (eds Ellison AM & Adamec L) 22–41 (Oxford University Press, 2018).

3 Hedrich, R. & Fukushima, K. On the origin of carnivory: molecular physiology and evolution of plants on an animal diet. Annual review of plant biology 72, 133–153 (2021).

4 Adamec, L., Matušíková, I. & Pavlovič, A. Recent ecophysiological, biochemical and evolutional insights into plant carnivory. Annals of Botany 128, 241–259 (2021).

5 Ellison, A. M. & Adamec, L. Carnivorous Plants: physiology, ecology, and evolution. (Oxford University Press, 2018).

6 Płachno, B. & Muravnik, L. E. Functional anatomy of carnivorous traps. (2018).

7 Leushkin, E. V. et al. The miniature genome of a carnivorous plant Genlisea aurea contains a low number of genes and short non-coding sequences. BMC genomics 14, 1–11 (2013).

8 Lan, T. et al. Long-read sequencing uncovers the adaptive topography of a carnivorous plant genome. Proceedings of the National Academy of Sciences 114, E4435–E4441 (2017).

9 Silva, S. R. et al. The terrestrial carnivorous plant Utricularia reniformis sheds light on environmental and life-form genome plasticity. International journal of molecular sciences 21, 3 (2019).

10 Vu, G. T. et al. Comparative genome analysis reveals divergent genome size evolution in a carnivorous plant genus. The Plant Genome 8, plantgenome2015.2004.0021 (2015).

11 Procko, C., Chory, J. & Pirro, S. The Genome Sequences of 17 Species of Carnivorous Plants. Biodiversity genomes (2023).

12 Saul, F. et al. Subgenome dominance shapes novel gene evolution in the decaploid pitcher plant Nepenthes gracilis. bioRxiv, 2023.2006. 2014.544965 (2023).

13 Kocáb, O. et al. Jasmonate-independent regulation of digestive enzyme activity in the carnivorous butterwort Pinguicula× Tina. Journal of Experimental Botany 71, 3749–3758 (2020).

14 Refulio-Rodriguez, N. F. & Olmstead, R. G. Phylogeny of lamiidae. American Journal of Botany 101, 287–299 (2014).

15 Zhang, C. et al. Asterid phylogenomics/phylotranscriptomics uncover morphological evolutionary histories and support phylogenetic placement for numerous whole-genome duplications. Molecular biology and evolution 37, 3188–3210 (2020).

16 Xu, W. Q. et al. Comparative genomics of figworts (Scrophularia, Scrophulariaceae), with implications for the evolution of Scrophularia and Lamiales. Journal of systematics and evolution 57, 55–65 (2019).

17 Initiative, O. T. P. T. One thousand plant transcriptomes and the phylogenomics of green plants. Nature 574, 679–685 (2019).

18 Liu, B. et al. Phylogenetic relationships of Cyrtandromoea and Wightia revisited: A new tribe in Phrymaceae and a new family in Lamiales. Journal of Systematics and Evolution 58, 1–17 (2020).

19 Veleba, A. et al. Genome size and genomic GC content evolution in the miniature genome-sized family Lentibulariaceae. New Phytologist 203, 22–28 (2014).

20 Guiglielmoni, N., Houtain, A., Derzelle, A., Van Doninck, K. & Flot, J.-F. Overcoming uncollapsed haplotypes in long-read assemblies of non-model organisms. BMC bioinformatics 22, 1–23 (2021).

21 Shimai, H., Setoguchi, H., Roberts, D. L. & Sun, M. Biogeographical patterns and speciation of the genus Pinguicula (Lentibulariaceae) inferred by phylogenetic analyses. PLoS One 16, e0252581 (2021).

22 Casper, S. J. & Stimper, R. Chromosome numbers in Pinguicula (Lentibulariaceae): survey, atlas, and taxonomic conclusions. Plant Systematics and Evolution 277, 21–60 (2009).

23 Ibarra-Laclette, E. et al. Architecture and evolution of a minute plant genome. Nature 498, 94–98 (2013).

24 Paysan-Lafosse, T. et al. InterPro in 2022. Nucleic acids research 51, D418–D427 (2023).

25 Jobson, R. W., Playford, J., Cameron, K. M. & Albert, V. A. Molecular phylogenetics of Lentibulariaceae inferred from plastid rps16 intron and trnL-F DNA sequences: implications for character evolution and biogeography. Systematic Botany, 157–171 (2003).

26 Jobson, R. W. & Albert, V. A. Molecular Rates Parallel Diversification Contrasts between Carnivorous Plant Sister Lineages. Cladistics 18, 127–136 (2002). 10.1006/clad.2001.0187

27 Ibarra-Laclette, E. et al. Transcriptomics and molecular evolutionary rate analysis of the bladderwort (Utricularia), a carnivorous plant with a minimal genome. BMC Plant Biology 11, 101 (2011). 10.1186/1471-2229-11-101

28 Arai, N., Ohno, Y., Jumyo, S., Hamaji, Y. & Ohyama, T. Organ-specific expression and epigenetic traits of genes encoding digestive enzymes in the lance-leaf sundew (Drosera adelae). Journal of Experimental Botany 72, 1946–1961 (2021).

29 *TAIR*: The Arabidopsis Information Resource (TAIR), <https://www.arabidopsis.org/download/index-auto.jsp%3Fdir%3D%252Fdownload_files%252FGenes%252FTAIR10_genome_release on www.arabidopsis.org, 05/01/2022> (2019).

30 Teshima, K. M. & Innan, H. The effect of gene conversion on the divergence between duplicated genes. Genetics 166, 1553–1560 (2004).

31 De Coster, W., D’Hert, S., Schultz, D. T., Cruts, M. & Van Broeckhoven, C. NanoPack: visualizing and processing long-read sequencing data. Bioinformatics 34, 2666–2669 (2018).

32 Andrews, S. (Babraham Bioinformatics, Babraham Institute, Cambridge, United Kingdom, 2010).

33 Li, H. Minimap and miniasm: fast mapping and de novo assembly for noisy long sequences. Bioinformatics 32, 2103–2110 (2016).

34 Li, H. Minimap2: pairwise alignment for nucleotide sequences. Bioinformatics 34, 3094–3100 (2018).

35 Vaser, R., Sović, I., Nagarajan, N. & Šikić, M. Fast and accurate de novo genome assembly from long uncorrected reads. Genome research 27, 737–746 (2017).

36 Walker, B. J. et al. Pilon: an integrated tool for comprehensive microbial variant detection and genome assembly improvement. PloS one 9, e112963 (2014).

37 Li, H. et al. The sequence alignment/map format and SAMtools. Bioinformatics 25, 2078–2079 (2009).

38. 38 Li, H. Aligning sequence reads, clone sequences and assembly contigs with BWA-MEM. *arXiv preprint arXiv:1303.*3997 (2013).

39 Roach, M. J., Schmidt, S. A. & Borneman, A. R. Purge Haplotigs: allelic contig reassignment for third-gen diploid genome assemblies. BMC bioinformatics 19, 1–10 (2018).

40 Lieberman-Aiden, E. et al. Comprehensive mapping of long-range interactions reveals folding principles of the human genome. science 326, 289–293 (2009).

41 Putnam, N. H. et al. Chromosome-scale shotgun assembly using an in vitro method for long-range linkage. Genome research 26, 342–350 (2016).

42. 42 Zaharia, M., et al. Faster and more accurate sequence alignment with SNAP. *arXiv preprint arXiv:1111.*5572 (2011).

43 Bolosky, W. J. et al. Fuzzy set intersection based paired-end short-read alignment. bioRxiv, 2021.2011. 2023.469039 (2021).

44 Alonge, M. et al. Automated assembly scaffolding using RagTag elevates a new tomato system for high-throughput genome editing. Genome biology 23, 1–19 (2022).

45 Gurevich, A., Saveliev, V., Vyahhi, N. & Tesler, G. QUAST: quality assessment tool for genome assemblies. Bioinformatics 29, 1072–1075 (2013).

46 Manni, M., Berkeley, M. R., Seppey, M., Simão, F. A. & Zdobnov, E. M. BUSCO update: novel and streamlined workflows along with broader and deeper phylogenetic coverage for scoring of eukaryotic, prokaryotic, and viral genomes. Molecular biology and evolution 38, 4647–4654 (2021).

47 Haas, B. J. et al. Automated eukaryotic gene structure annotation using EVidenceModeler and the Program to Assemble Spliced Alignments. Genome biology 9, 1–22 (2008).

48 Haas, B. J., Zeng, Q., Pearson, M. D., Cuomo, C. A. & Wortman, J. R. Approaches to fungal genome annotation. Mycology 2, 118–141 (2011).

49 Bolger, A. M., Lohse, M. & Usadel, B. Trimmomatic: a flexible trimmer for Illumina sequence data. Bioinformatics 30, 2114–2120 (2014).

50 Smit, A. & Hubley, R. RepeatModeler Open-1.0, <http://www.repeatmasker.org> (2008-2015).

51 Smit, A., Hubley, R. & Green, P. RepeatMasker Open-4.0, <http://www.repeatmasker.org> (2013-2015).

52 Price, A. L., Jones, N. C. & Pevzner, P. A. De novo identification of repeat families in large genomes. Bioinformatics 21, i351–i358 (2005).

53 Bao, Z. & Eddy, S. R. Automated de novo identification of repeat sequence families in sequenced genomes. Genome research 12, 1269–1276 (2002).

54 Ellinghaus, D., Kurtz, S. & Willhoeft, U. LTRharvest, an efficient and flexible software for de novo detection of LTR retrotransposons. BMC bioinformatics 9, 1–14 (2008).

55 Ou, S. & Jiang, N. LTR_retriever: a highly accurate and sensitive program for identification of long terminal repeat retrotransposons. Plant physiology 176, 1410–1422 (2018).

56 Grabherr, M. G. et al. Full-length transcriptome assembly from RNA-Seq data without a reference genome. Nature biotechnology 29, 644–652 (2011).

57 Haas, B. J. et al. De novo transcript sequence reconstruction from RNA-seq using the Trinity platform for reference generation and analysis. Nature protocols 8, 1494–1512 (2013).

58 Kim, D., Paggi, J. M., Park, C., Bennett, C. & Salzberg, S. L. Graph-based genome alignment and genotyping with HISAT2 and HISAT-genotype. Nature biotechnology 37, 907–915 (2019).

59 Pertea, M. et al. StringTie enables improved reconstruction of a transcriptome from RNA-seq reads. Nature biotechnology 33, 290–295 (2015).

60 Dobin, A. et al. STAR: ultrafast universal RNA-seq aligner. Bioinformatics 29, 15–21 (2013).

61 Robertson, G. et al. De novo assembly and analysis of RNA-seq data. Nature methods 7, 909–912 (2010).

62 Gilbert, D. Genomes built from mRNA-seq not genome DNA. (2013).

63 Haas, B. J. et al. Improving the Arabidopsis genome annotation using maximal transcript alignment assemblies. Nucleic acids research 31, 5654–5666 (2003).

64 Wang, M. & Kong, L. pblat: a multithread blat algorithm speeding up aligning sequences to genomes. BMC bioinformatics 20, 1–4 (2019).

65 Wu, T. D. & Watanabe, C. K. GMAP: a genomic mapping and alignment program for mRNA and EST sequences. Bioinformatics 21, 1859–1875 (2005).

66 Keilwagen, J. et al. Using intron position conservation for homology-based gene prediction. Nucleic Acids Research 44, e89 (2016).

67 Keilwagen, J., Hartung, F. & Grau, J. in Gene prediction: Methods and protocols Vol. 1962 Methods in Molecular Biology (ed Martin Kollmar) Ch. 8, 161–177 (Springer Science+Business Media, LLC, part of Springer Nature, 2019).

68 Hellsten, U. et al. Fine-scale variation in meiotic recombination in Mimulus inferred from population shotgun sequencing. Proceedings of the National Academy of Sciences 110, 19478–19482 (2013).

69 Zhao, D. et al. A chromosomal-scale genome assembly of Tectona grandis reveals the importance of tandem gene duplication and enables discovery of genes in natural product biosynthetic pathways. Gigascience 8, giz005 (2019).

70 Wang, L. et al. Genome sequencing of the high oil crop sesame provides insight into oil biosynthesis. Genome biology 15, 1–13 (2014).

71 Hamilton, J. P. et al. Generation of a chromosome-scale genome assembly of the insect-repellent terpenoid-producing Lamiaceae species, Callicarpa americana. GigaScience 9, giaa093 (2020).

72 Lomsadze, A., Ter-Hovhannisyan, V., Chernoff, Y. O. & Borodovsky, M. Gene identification in novel eukaryotic genomes by self-training algorithm. Nucleic acids research 33, 6494–6506 (2005).

73 Hoff, K. J., Lange, S., Lomsadze, A., Borodovsky, M. & Stanke, M. BRAKER1: unsupervised RNA-Seq-based genome annotation with GeneMark-ET and AUGUSTUS. Bioinformatics 32, 767–769 (2016).

74 Hoff, K. J., Lomsadze, A., Borodovsky, M. & Stanke, M. Whole-genome annotation with BRAKER. Gene prediction: methods and protocols, 65–95 (2019).

75 Brůna, T., Hoff, K. J., Lomsadze, A., Stanke, M. & Borodovsky, M. BRAKER2: automatic eukaryotic genome annotation with GeneMark-EP+ and AUGUSTUS supported by a protein database. NAR genomics and bioinformatics 3, lqaa108 (2021).

76 Barnett, D. W., Garrison, E. K., Quinlan, A. R., Strömberg, M. P. & Marth, G. T. BamTools: a C++ API and toolkit for analyzing and managing BAM files. Bioinformatics 27, 1691–1692 (2011).

77 Stanke, M., Diekhans, M., Baertsch, R. & Haussler, D. Using native and syntenically mapped cDNA alignments to improve de novo gene finding. Bioinformatics 24, 637–644 (2008).

78 Stanke, M., Schöffmann, O., Morgenstern, B. & Waack, S. Gene prediction in eukaryotes with a generalized hidden Markov model that uses hints from external sources. BMC bioinformatics 7, 1–11 (2006).

79 Nakamura, T., Yamada, K. D., Tomii, K. & Katoh, K. Parallelization of MAFFT for large-scale multiple sequence alignments. Bioinformatics 34, 2490–2492 (2018).

80 Capella-Gutiérrez, S., Silla-Martínez, J. M. & Gabaldón, T. trimAl: a tool for automated alignment trimming in large-scale phylogenetic analyses. Bioinformatics 25, 1972–1973 (2009).

81 Nguyen, L.-T., Schmidt, H. A., Von Haeseler, A. & Minh, B. Q. IQ-TREE: a fast and effective stochastic algorithm for estimating maximum-likelihood phylogenies. Molecular biology and evolution 32, 268–274 (2015).

82 Hoang, D. T., Chernomor, O., Von Haeseler, A., Minh, B. Q. & Vinh, L. S. UFBoot2: improving the ultrafast bootstrap approximation. Molecular biology and evolution 35, 518–522 (2018).

83 Kalyaanamoorthy, S., Minh, B. Q., Wong, T. K., von Haeseler, A. & Jermiin, L. S. ModelFinder: fast model selection for accurate phylogenetic estimates. Nature methods 14, 587 (2017).

84 Zhang, C., Rabiee, M., Sayyari, E. & Mirarab, S. ASTRAL-III: polynomial time species tree reconstruction from partially resolved gene trees. BMC bioinformatics 19, 15–30 (2018).

85 Lyons, E. & Freeling, M. How to usefully compare homologous plant genes and chromosomes as DNA sequences. The Plant Journal 53, 661–673 (2008).

86 Haug-Baltzell, A., Stephens, S. A., Davey, S., Scheidegger, C. E. & Lyons, E. SynMap2 and SynMap3D: web-based whole-genome synteny browsers. Bioinformatics 33, 2197–2198 (2017).

87 Lyons, E., Freeling, M., Kustu, S. & Inwood, W. Using genomic sequencing for classical genetics in E. coli K12. PloS one 6, e16717 (2011).

88 Jaillon, O. et al. The grapevine genome sequence suggests ancestral hexaploidization in major angiosperm phyla. Nature 449, 463–467 (2007).

89 Murat, F. et al. Karyotype and gene order evolution from reconstructed extinct ancestors highlight contrasts in genome plasticity of modern rosid crops. Genome biology and evolution 7, 735–749 (2015).

90 Lyons, E., Pedersen, B., Kane, J. & Freeling, M. The value of nonmodel genomes and an example using SynMap within CoGe to dissect the hexaploidy that predates the rosids. Tropical Plant Biology 1, 181–190 (2008).

91 Joyce, B. L. et al. FractBias: a graphical tool for assessing fractionation bias following polyploidy. Bioinformatics 33, 552–554 (2017).

92 Tang, H. et al. Screening synteny blocks in pairwise genome comparisons through integer programming. BMC bioinformatics 12, 1–11 (2011).

93 Team”, R. C. R: A language and environment for statistical computing. (2013).

94 Wickham, H. et al. Welcome to the Tidyverse. Journal of open source software 4, 1686 (2019).

95 Hadley, W. Ggplot2: Elegrant graphics for data analysis. (Springer, 2016).

96 Neuwirth, E. & Neuwirth, M. E. Package ‘RColorBrewer’. ColorBrewer Palettes (2014).

97 Wilke, C. O. Ridgeline Plots in ‘ggplot2’[R Package Ggridges Version 0.5. 3]. January. https://cran.r-project.org/web/packages/ggridges/index.html (2021).

98. 98 Aphalo, P. (2020).

99 Morgan, M., Obenchain, V., Hester, J. & Pagès, H. SummarizedExperiment: A container (S4 class) for matrix-like assays. R package version 1.36.0, <https://bioconductor.org/packages/SummarizedExperiment.> (2024).

100 Lyons, E. et al. Finding and comparing syntenic regions among Arabidopsis and the outgroups papaya, poplar, and grape: CoGe with rosids. Plant physiology 148, 1772–1781 (2008).

101 Tang, H. et al. Synteny and collinearity in plant genomes. Science 320, 486–488 (2008).

102 Dainat, J. AGAT: Another Gff Analysis Toolkit to handle annotations in any GTF/GFF format (version 0.9.1), <https://zenodo.org/records/6488306> (2022).

103 Camacho, C. et al. BLAST+: architecture and applications. BMC bioinformatics 10, 1–9 (2009).

104 Denoeud, F. et al. The coffee genome provides insight into the convergent evolution of caffeine biosynthesis. science 345, 1181–1184 (2014).

105 Xu, Z. et al. Tandem gene duplications drive divergent evolution of caffeine and crocin biosynthetic pathways in plants. BMC biology 18, 1–14 (2020).

106 Krzywinski, M. et al. Circos: an information aesthetic for comparative genomics. Genome research 19, 1639–1645 (2009).

