## Supplementary figures for "Genomic Insights into the Evolution of Carnivory in the Giant Butterwort, *Pinguicula gigantea*: Chromosome-Level Assembly and Phylogenomic Analysis"

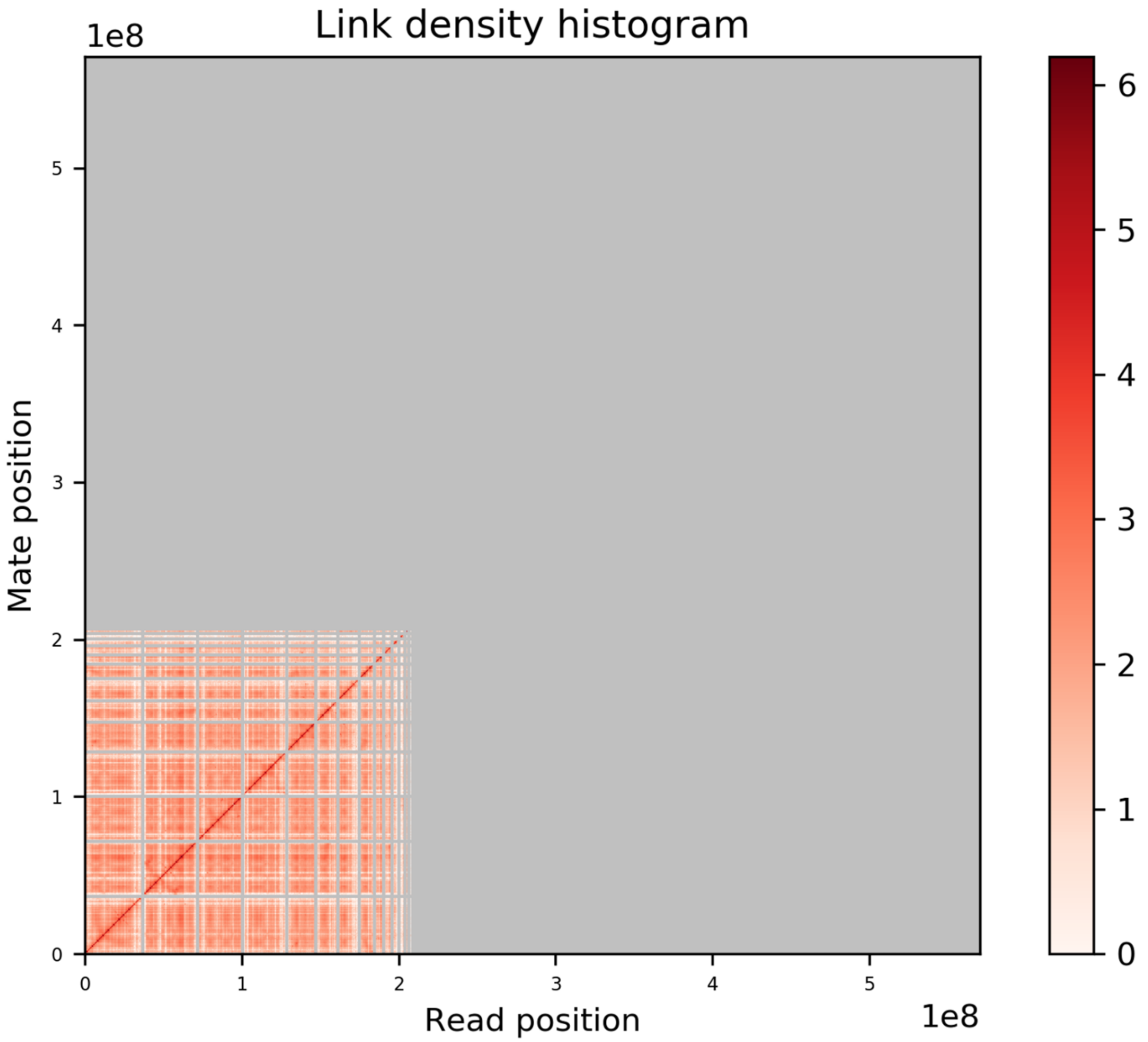


Figure S1: Hi-C heat map for *Pinguicula grandiflora*. Darker red indicates a stronger signal. Read position on the Left and bottom.

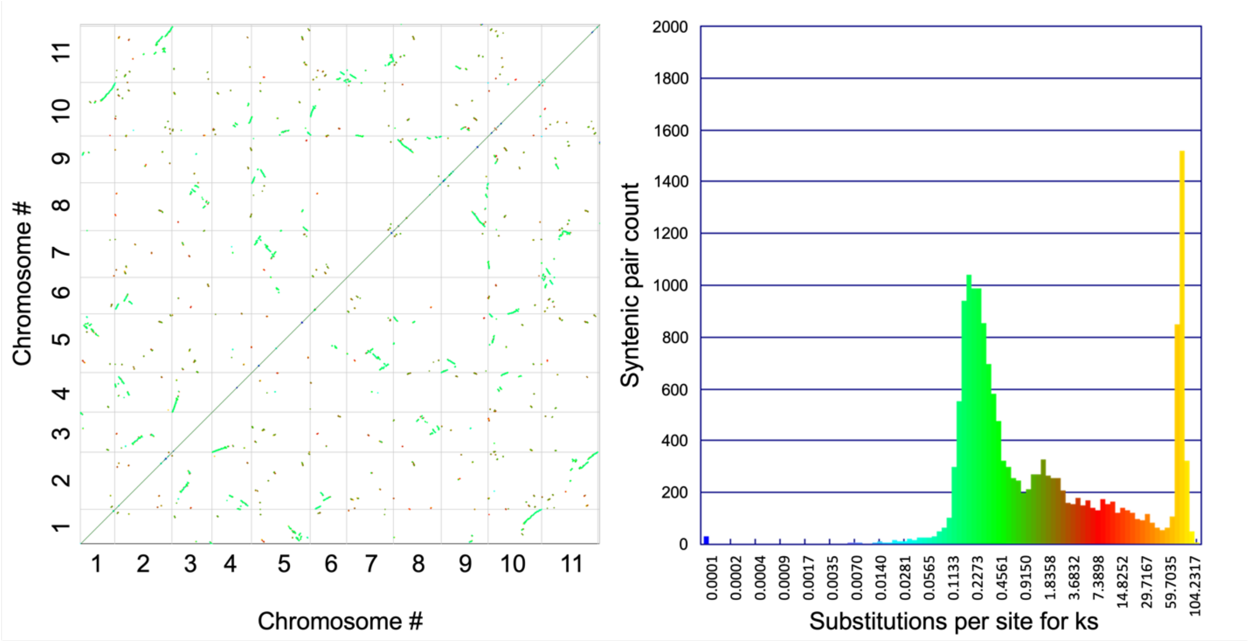


Figure S2: *Pinguicula gigantea* Hi-C assembly syntenic dot plot and syntenic gene-pair Ks histogram. (left) Each dot represents a syntenic paralogous gene pair. *Pinguicula gigantea* genome is along the X and Y-axis. (right) Histogram of synonymous substitution rate between syntenic paralogous gene pairs. Syntenic gene pair count is on the X-axis and Ks is on the X-axis. The color of the dot in the dot plot (left) corresponds with the Ks value on the Ks histogram (right).

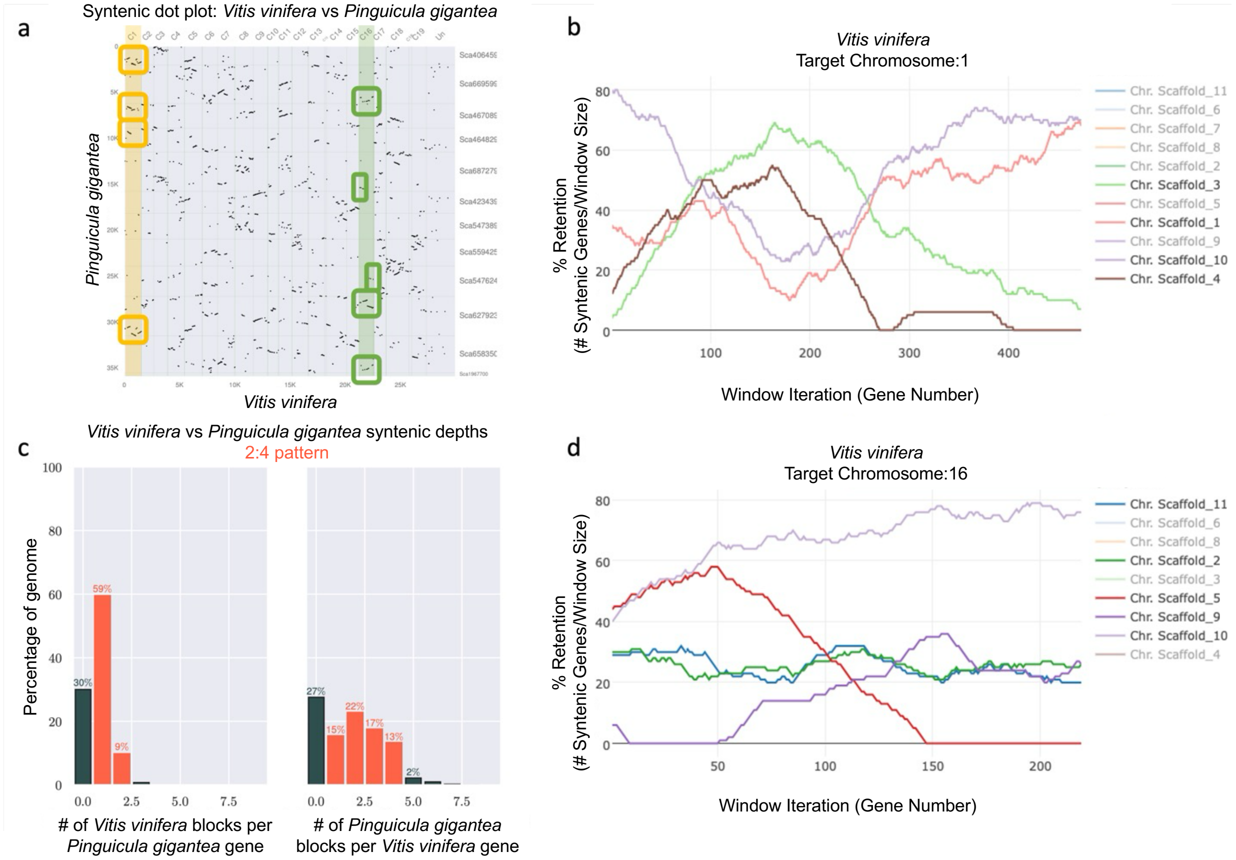


Figure S3: Comparative ploidy level between *Pinguicula gigantea* and *Vitis vinifera*. (a) Each dot represents a syntenic orthologous gene pair between *Pinguicula gigantea* on the Y-axis and *Vitis vinifera* on the X-axis*. Vitis vinifera* chromosomes 1 and 16 are highlighted in yellow and green, respectively. (b) Fraction bias plot with *Vitis vinifera* chromosome 1 on the X-axis and percent gene retention on the Y-axis. Different color lines represent different chromosomes from *Pinguicula gigantea*. (c) Syntenic depth histogram. The left histogram is number of *Vitis vinifera* blocks per *Pinguicula gigantea* gene. The right histogram is number of *Pinguicula gigantea* blocks per *Vitis vinifera* gene. (d) Fraction bias plot with *Vitis vinifera* chromosome 16 on the X-axis and percent gene retention on the Y-axis. Different color lines represent different chromosomes from *Pinguicula gigantea*.

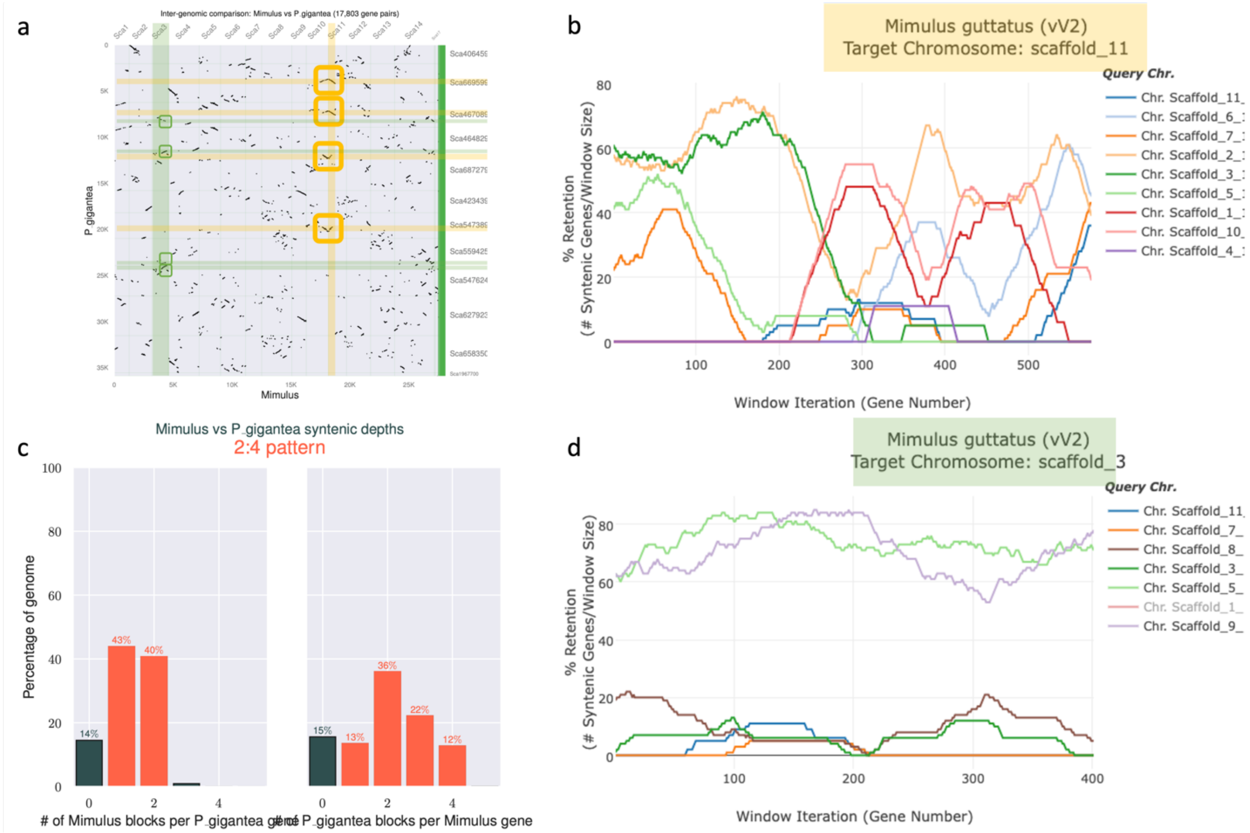


Figure S4: Comparative ploidy level between *Pinguicula gigantea* and *Mimulus guttatus*. (a) Each dot represents a syntenic orthologous gene pair between *Pinguicula gigantea* on the Y-axis and *Mimulus guttatus* on the X-axis*. Mimulus guttatus* chromosome 11 and 3 are highlighted in yellow and green, respectively. (b) Fraction bias plot with *Mimulus guttatus* chromosome 11 on the X-axis and percent gene retention on the Y-axis. Different color lines represent different chromosomes from *Pinguicula gigantea*. (c) Syntenic depth histogram. The left histogram is number of *Mimulus guttatus* blocks per *Pinguicula gigantea* gene. The right histogram is number of *Pinguicula gigantea* blocks per *Mimulus guttatus* gene. (d) Fraction bias plot with *Mimulus guttatus* chromosome 3 on the X-axis and percent gene retention on the Y-axis. Different color lines represent different chromosomes from *Pinguicula gigantea*.

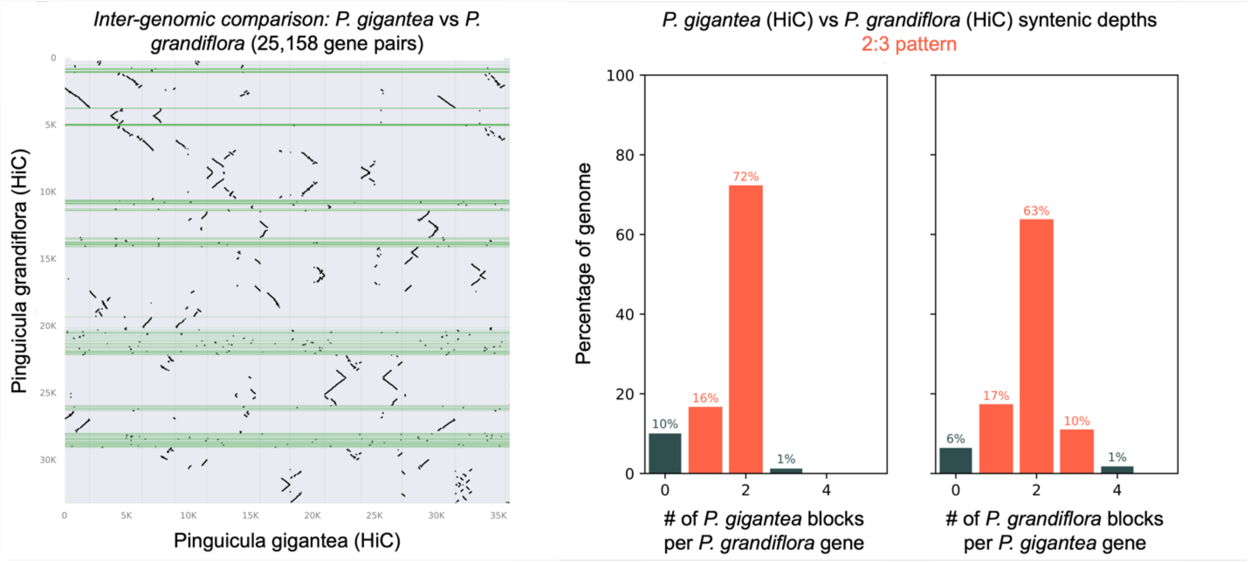


Figure S5: Comparative ploidy level between the *Pinguicula grandiflora* Hi-C assembly and the *Pinguicula gigantea* Hi-C assembly. (left) Each dot represents a syntenic orthologous gene pair between *Pinguicula grandiflora* on the Y-axis and *Pinguicula gigantea* on the X-axis. (right) Syntenic depth histogram. The left histogram is number of *Pinguicula gigantea* blocks per *Pinguicula grandiflora* gene. The right histogram is number of *Pinguicula grandiflora* blocks per *Pinguicula gigantea* gene.

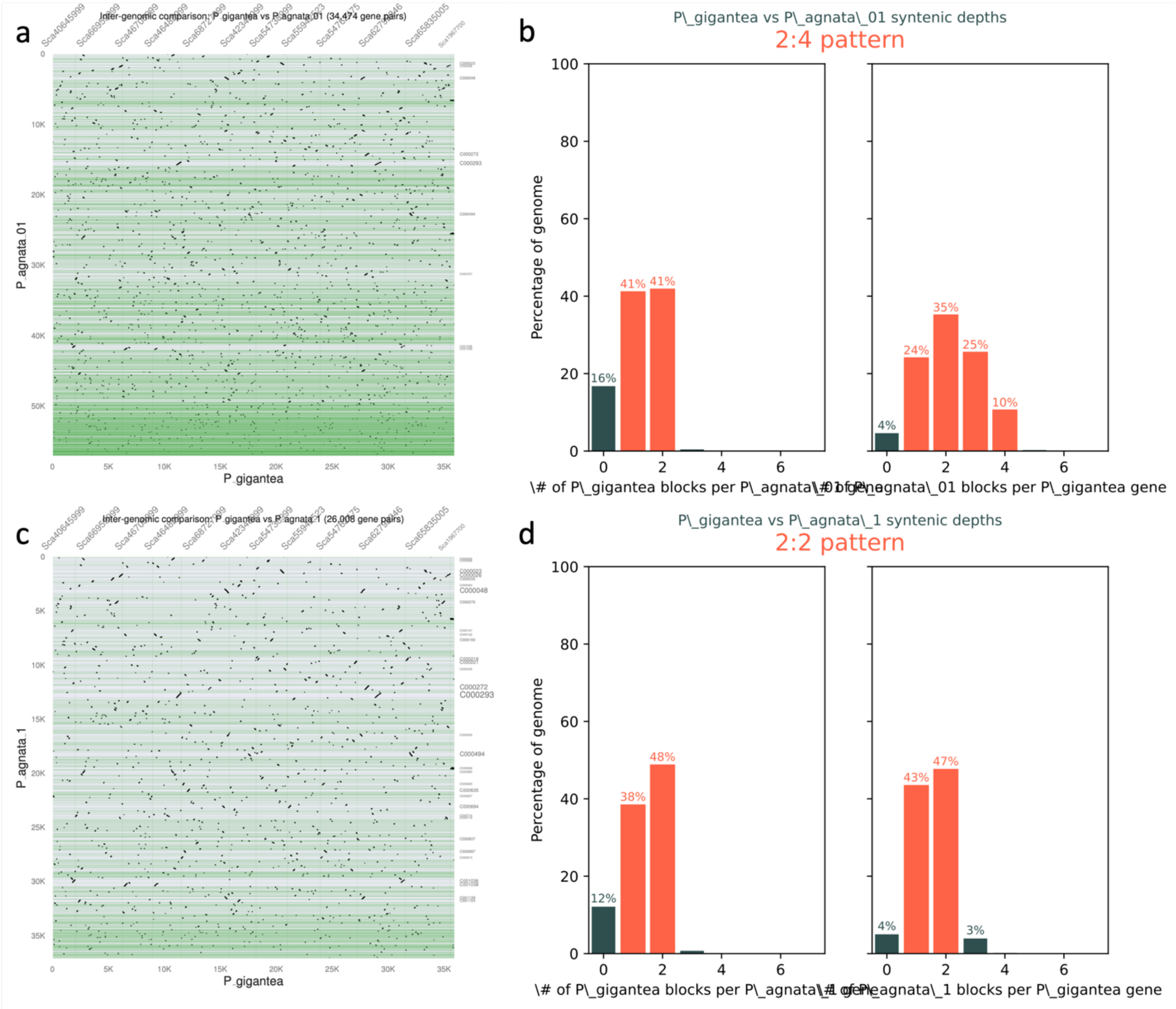


Figure S6: *Pinguicula agnata* long-read assembly vs *Pinguicula gigantea* Hi-C assembly syntenic dot plot and syntenic depth. (a and b) *Pinguicula agnata* primary assembly before PurgeHaplotigs. (c and d) *Pinguicula agnata* assembly after PurgeHaplotigs. (left) Each dot represents a syntenic orthologous gene pair between *Pinguicula agnata* and *Pinguicula gigantea*. *Pinguicula agnata* genome is along the X-axis and *Pinguicula gigantea* is along the Y-axis. (right) Syntenic depth histogram. The left histogram is number of *Pinguicula gigantea* blocks per *Pinguicula agnata* gene. The right histogram is number of *Pinguicula agnata* blocks per *Pinguicula gigantea* gene.

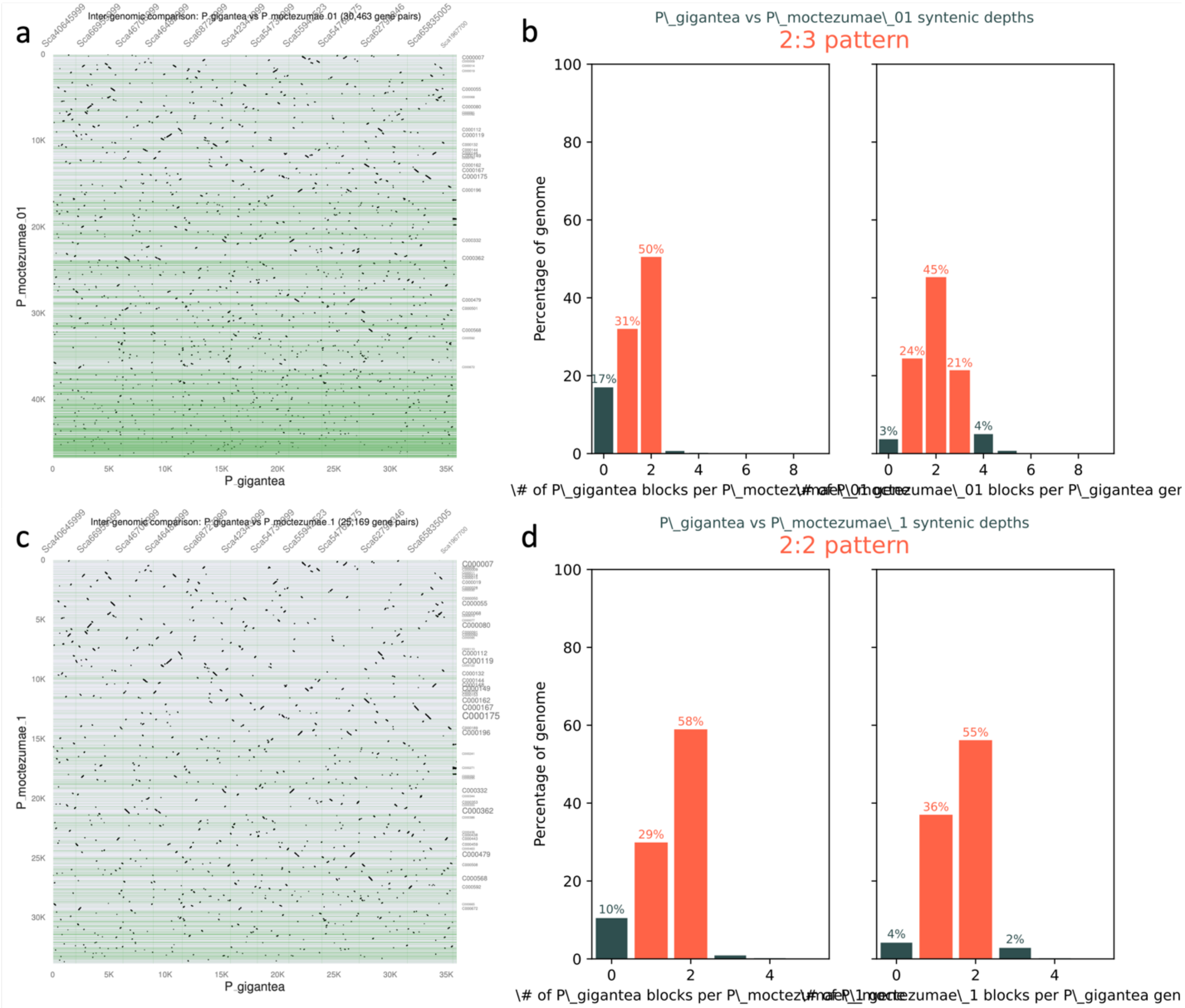


Figure S7: *Pinguicula moctezumae* long-read assembly vs *Pinguicula gigantea* Hi-C assembly syntenic dot plot and syntenic depth. (a and b) *Pinguicula moctezumae* primary assembly before PurgeHaplotigs. (c and d) *Pinguicula moctezumae* assembly after PurgeHaplotigs. (left) Each dot represents a syntenic orthologous gene pair between *Pinguicula moctezumae* and *Pinguicula gigantea*. *Pinguicula moctezumae* genome is along the X-axis and *Pinguicula gigantea* is along the Y-axis. (right) Syntenic depth histogram. The left histogram is number of *Pinguicula gigantea* blocks per *Pinguicula moctezumae* gene. The right histogram is number of *Pinguicula moctezumae* blocks per *Pinguicula gigantea* gene.

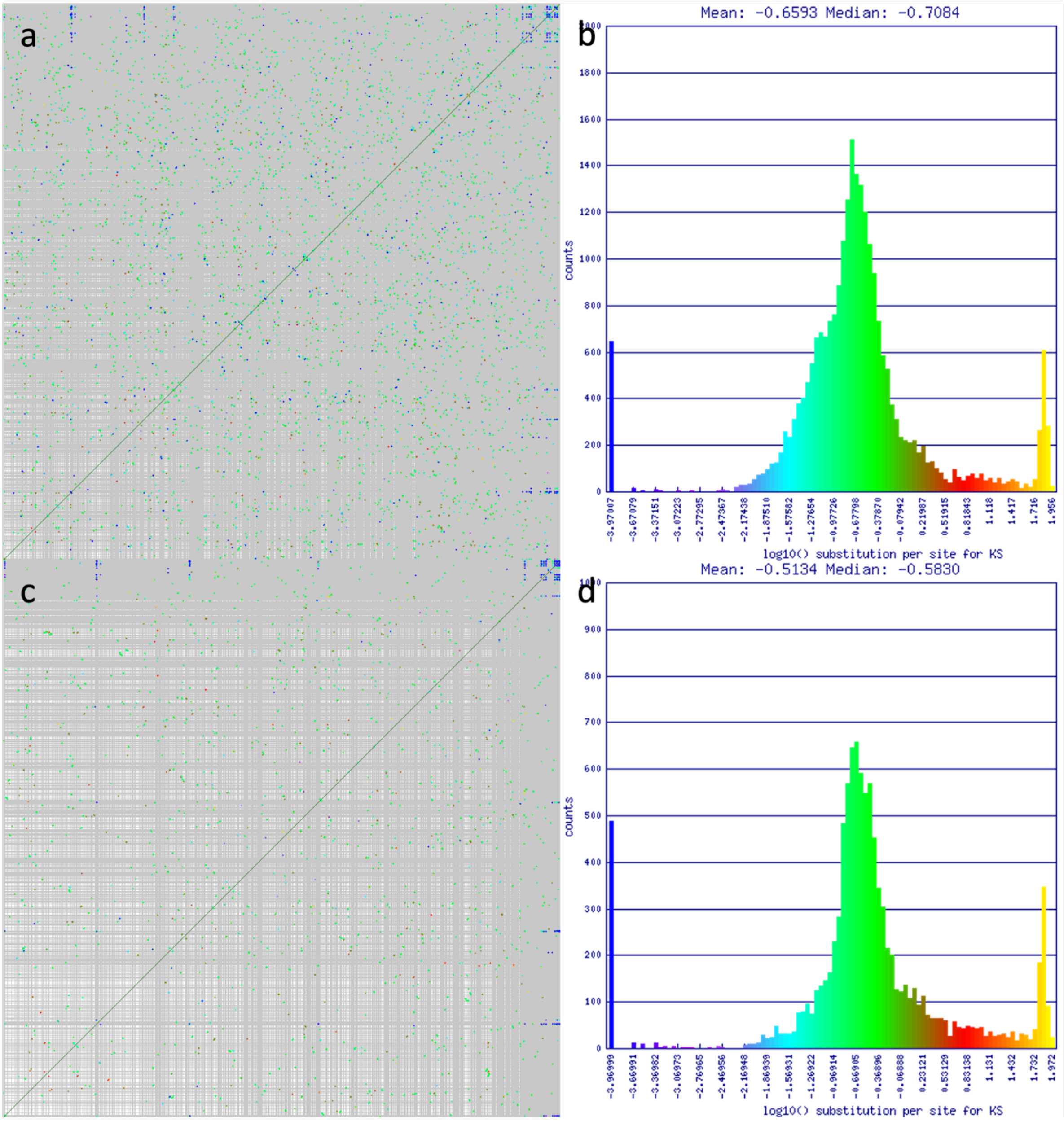

Figure S8: *Pinguicula agnata* long-read assembly syntenic dot plot and syntenic gene-pair Ks histogram. (a and b) *Pinguicula agnata* primary assembly before PurgeHaplotigs. (c and d) *Pinguicula agnata* assembly after PurgeHaplotigs. (left) Each dot represents a syntenic paralogous gene pair. *Pinguicula agnata* genome is along the X and Y-axis. (right) Histogram of synonymous substitution rate between syntenic paralogous gene pairs. Syntenic gene pair count is on the X-axis and Ks is on the X-axis. The color of the dot in the dot plot (left) corresponds with the Ks value on the Ks histogram (right).

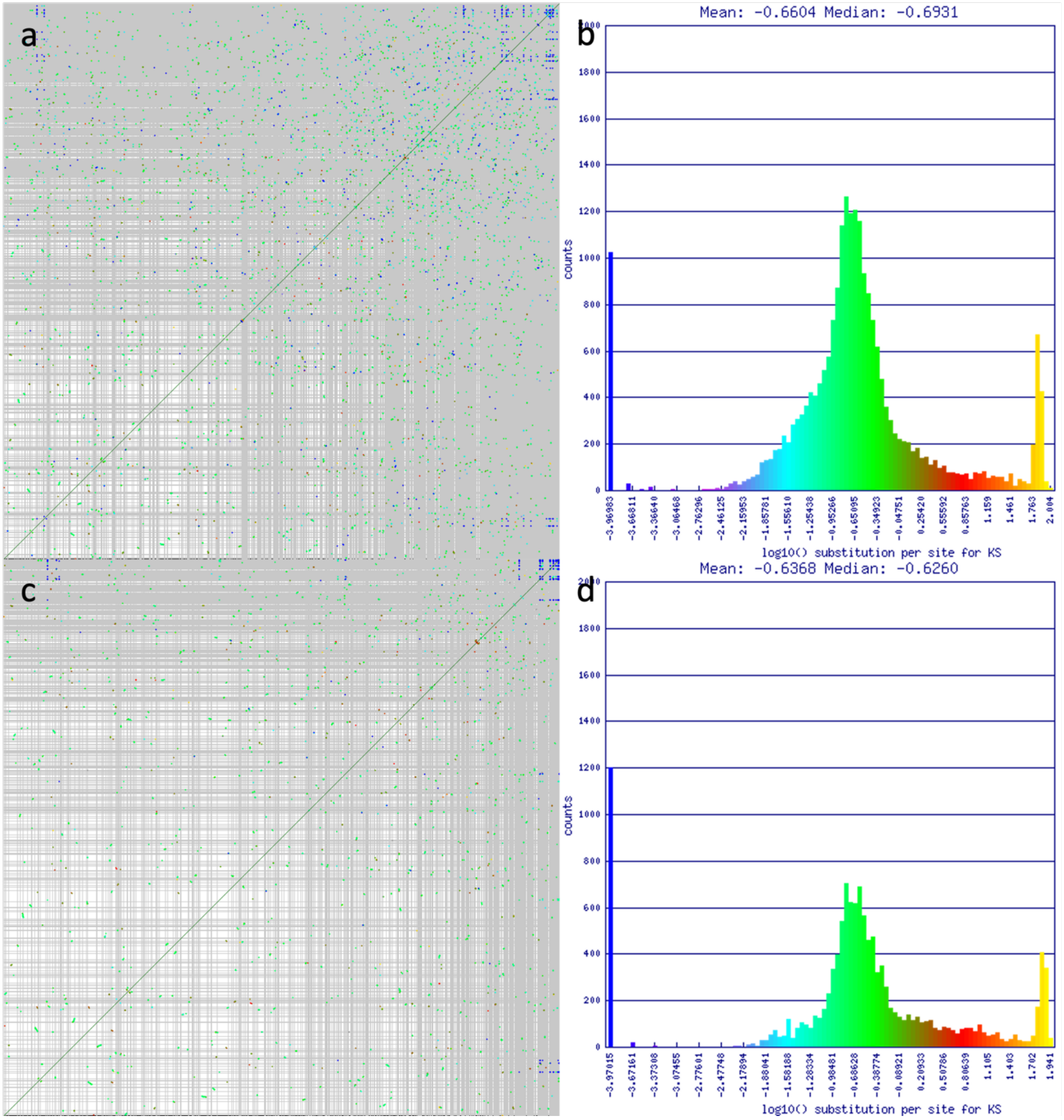


Figure S9: *Pinguicula moctezumae* long-read assembly syntenic dot plot and syntenic gene-pair Ks histogram. (a and b) *Pinguicula moctezumae* primary assembly before PurgeHaplotigs. (c and d) *Pinguicula moctezumae* assembly after PurgeHaplotigs. (left) Each dot represents a syntenic paralogous gene pair. *Pinguicula moctezumae* genome is along the X and Y-axis. (right) Histogram of synonymous substitution rate between syntenic paralogous gene pairs. Syntenic gene pair count is on the X-axis and Ks is on the X-axis. The color of the dot in the dot plot (left) corresponds with the Ks value on the Ks histogram (right).

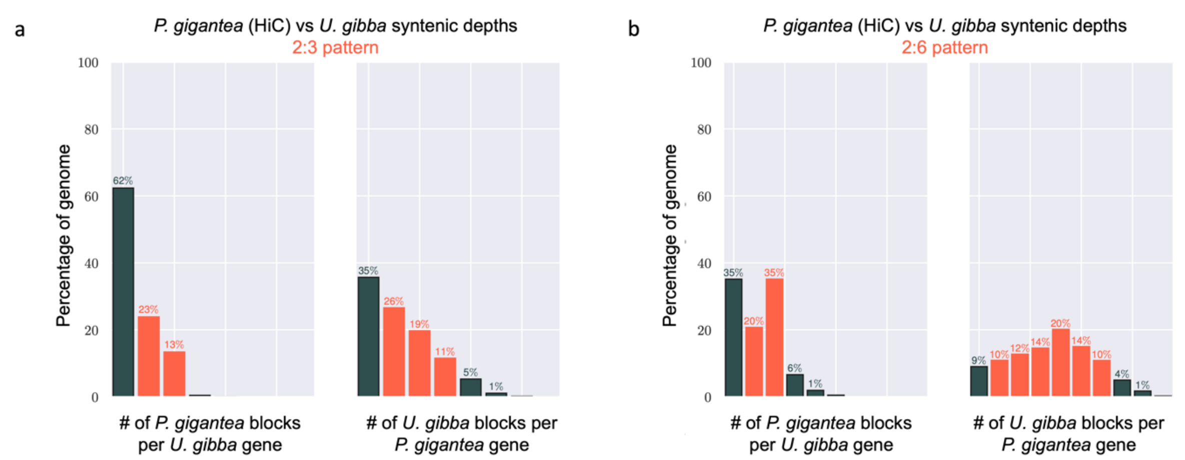


Figure S10: Comparative ploidy levels between *Pinguicula gigantea* and *Utricularia gibba*. (a) Syntenic depth histogram generated by searching for five syntenic genes within twenty nearby genes. The left histogram is number of *Pinguicula gigantea* blocks per *Utricularia gibba* gene. The right histogram is number of *Utricularia gibba* blocks per *Pinguicula gigantea* gene. (b) Syntenic depth histogram generated by searching for five syntenic genes within fifty nearby genes. The left histogram is number of *Pinguicula gigantea* blocks per *Utricularia gibba* gene. The right histogram is number of *Utricularia gibba* blocks per *Pinguicula gigantea* gene.

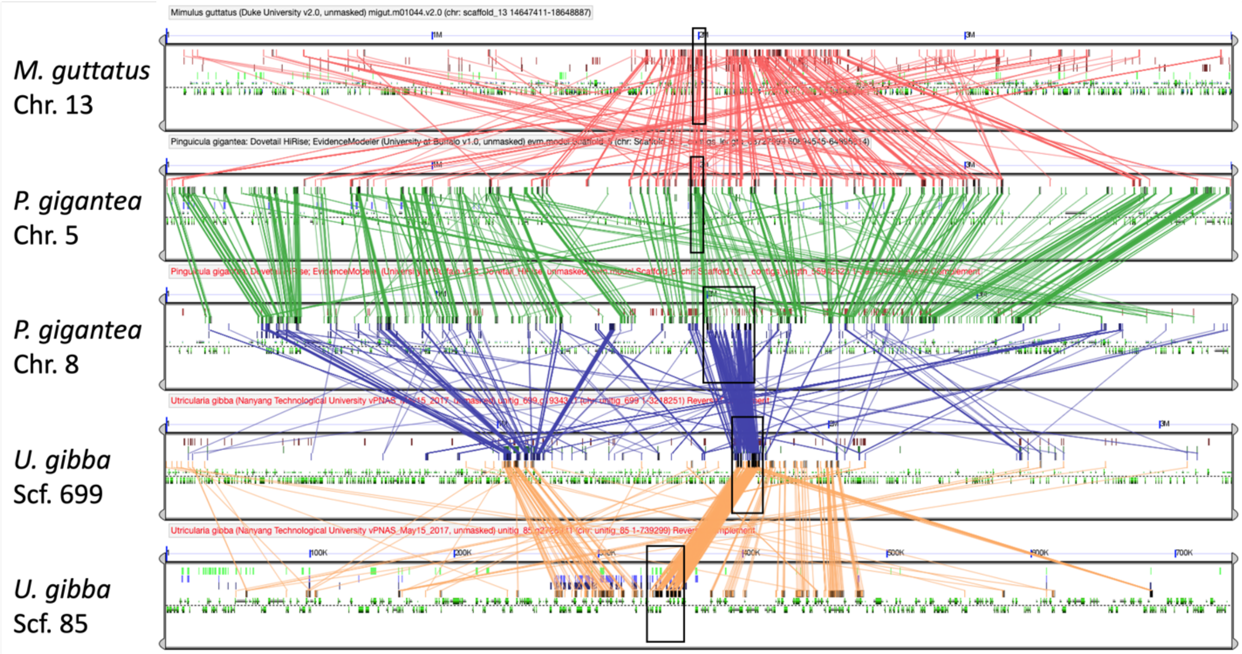


Figure S11: Cysteine protease tandem array in *Pinguicula gigantea* and homologous regions in *Mimulus guttatus* and *Utricularia gibba*. Syntenic blocks between *Mimulus guttatus* scaffold 13, *Pinguicula gigantea* chromosome 5, *Pinguicula gigantea* chromosome 8, *Utricularia gibba* scaffold 699, and *Utricularia gibba* scaffold 85. The tandem arrays on *Pinguicula gigantea* chromosome 8, *Utricularia gibba* scaffold 699, and *Utricularia gibba* scaffold 85 and the single cysteine protease genes on *Mimulus guttatus* scaffold 13 and *Pinguicula gigantea* chromosome 5 are marked with black rectangles. CDS are represented by green rectangles. Colored lines connect homologous gene pairs.

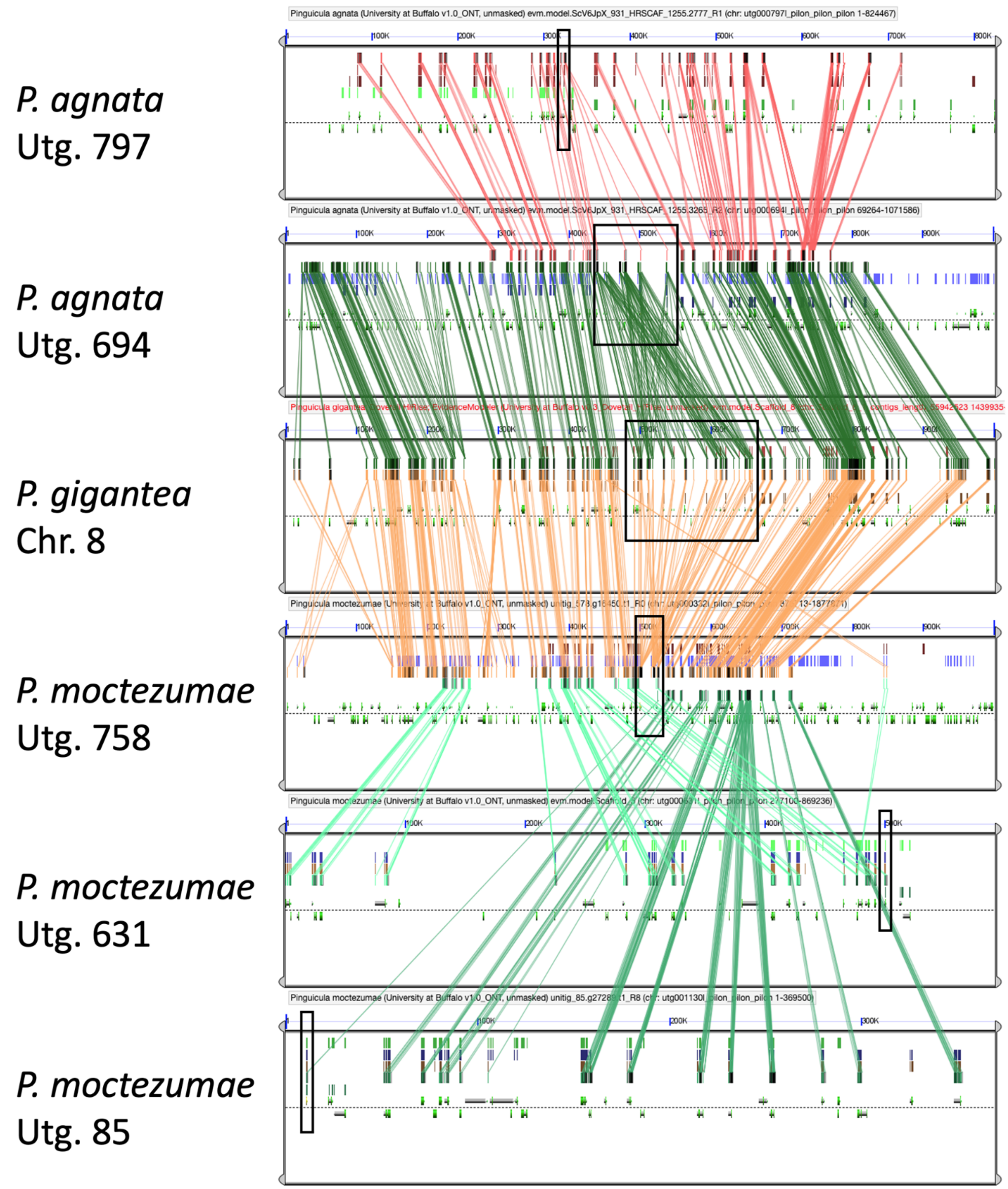


Figure S12: Cysteine protease tandem array in *Pinguicula gigantea* and homologous regions in *Pinguicula agnata* and *Pinguicula moctezumae*. Syntenic blocks between *Pinguicula agnata* unitigs 797 and 694, *Pinguicula gigantea* chromosome 8, and *Pinguicula moctezumae* unitigs 758, 631, and 85. The tandem arrays on *Pinguicula agnata* unitig 694, *Pinguicula gigantea* chromosome 8, and *Pinguicula moctezumae* unitig 758 and the single cysteine protease genes on *Pinguicula agnata* unitig 797 and

*Pinguicula moctezumae* unitig 85 are marked with black rectangles. CDS are represented by green rectangles. Colored lines connect homologous gene pairs.

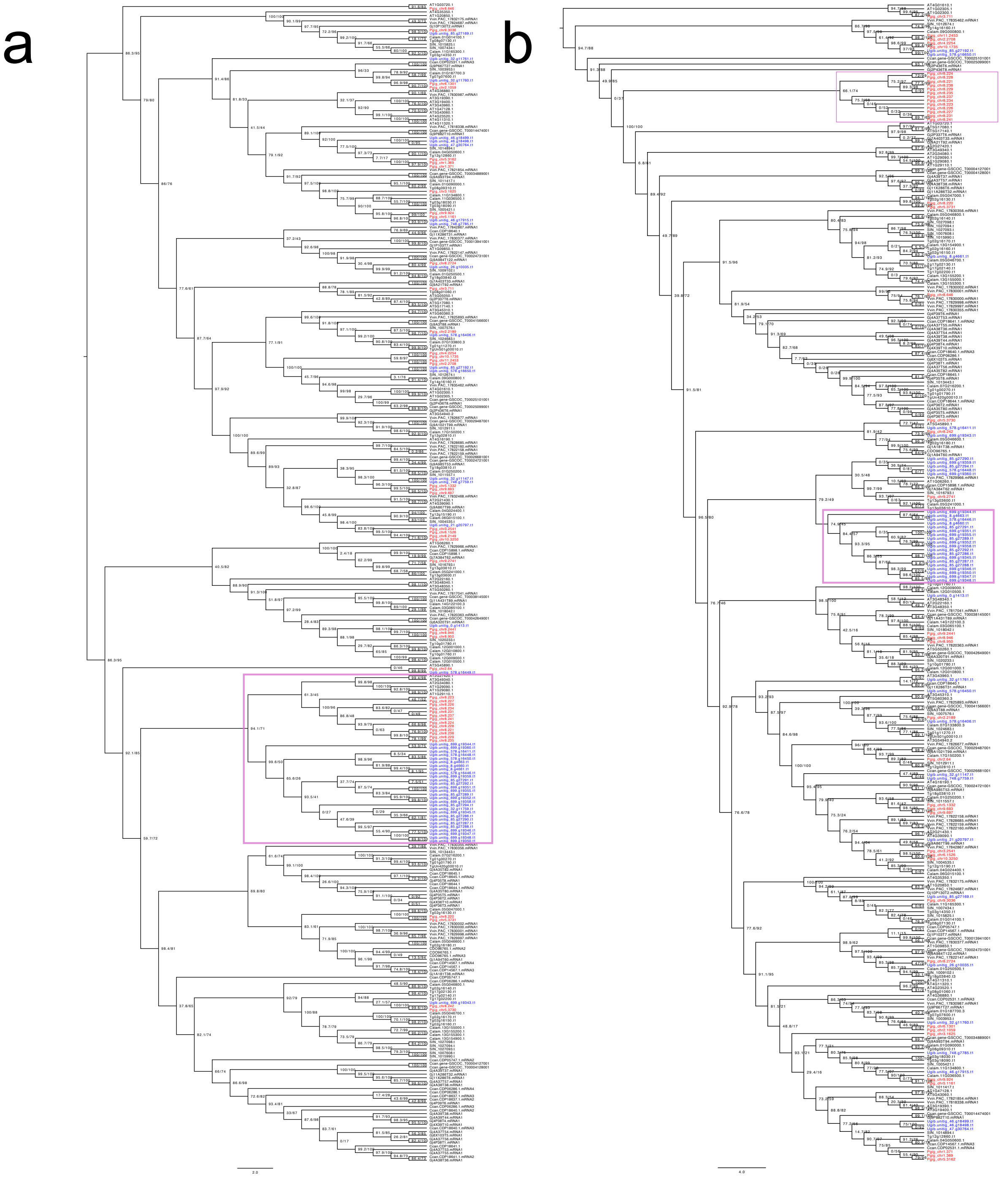


Figure S13: Cysteine protease CDS and protein trees. (a) CDS sequences. (b) Protein sequences. All clades containing *Pinguicula gigantea* sequences with three or more tandem genes are highlighted and zoomed in. Branch support is the ultrafast bootstrap replicate score. *Utricularia gibba* sequences are in blue and *Pinguicula gigantea* sequences are in red. Tandem duplicates of 3 or more that group together on the tree are boxed in purple.

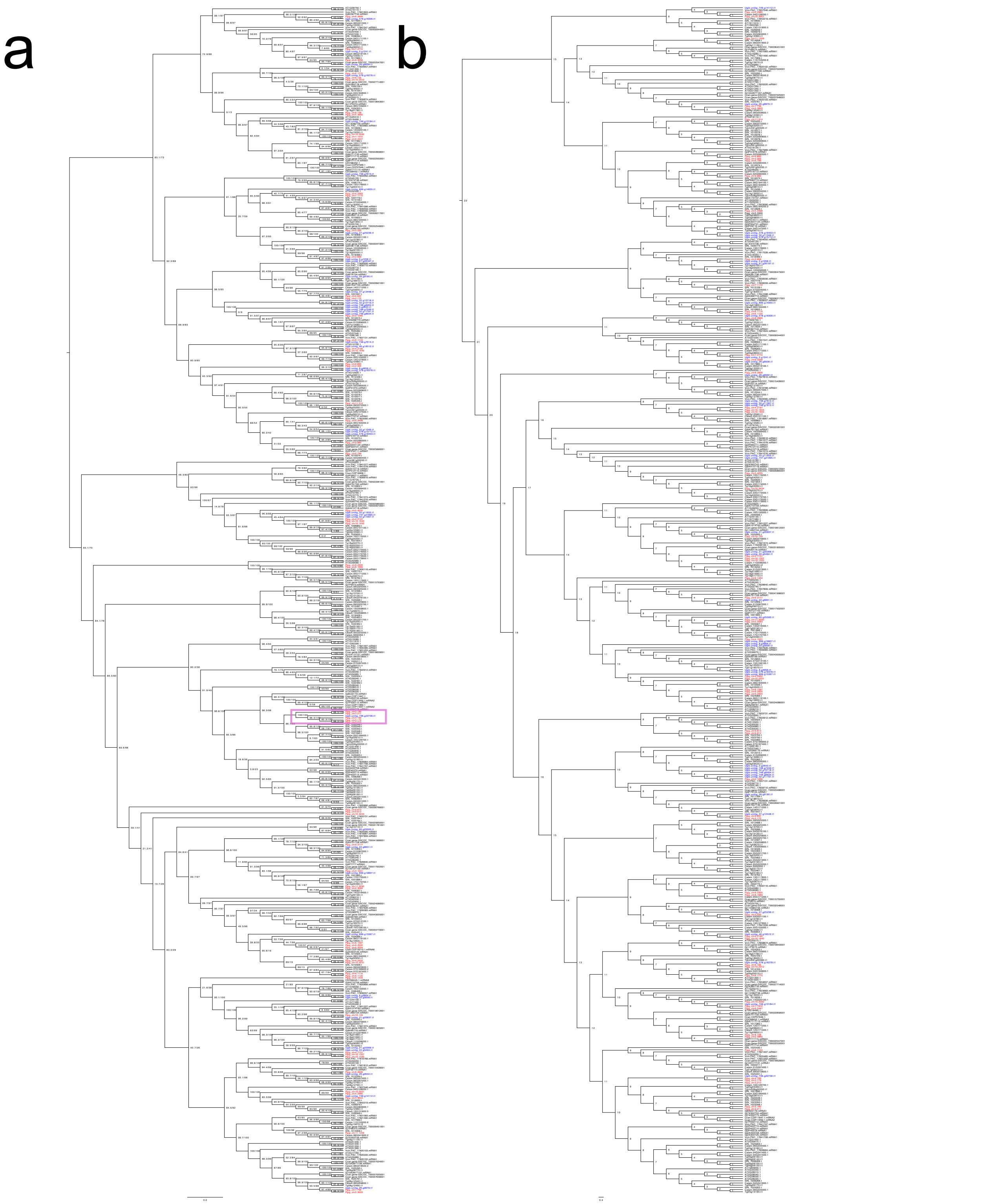


Figure S14: Aspartic protease CDS and protein trees. (a) CDS sequences. (b) Protein sequences. All clades containing *Pinguicula gigantea* sequences with three or more tandem genes are highlighted and zoomed in. Branch support is the ultrafast bootstrap replicate score. *Utricularia gibba* sequences are in blue and *Pinguicula gigantea* sequences are in red. Tandem duplicates of 3 or more that group together on the tree are boxed in purple.

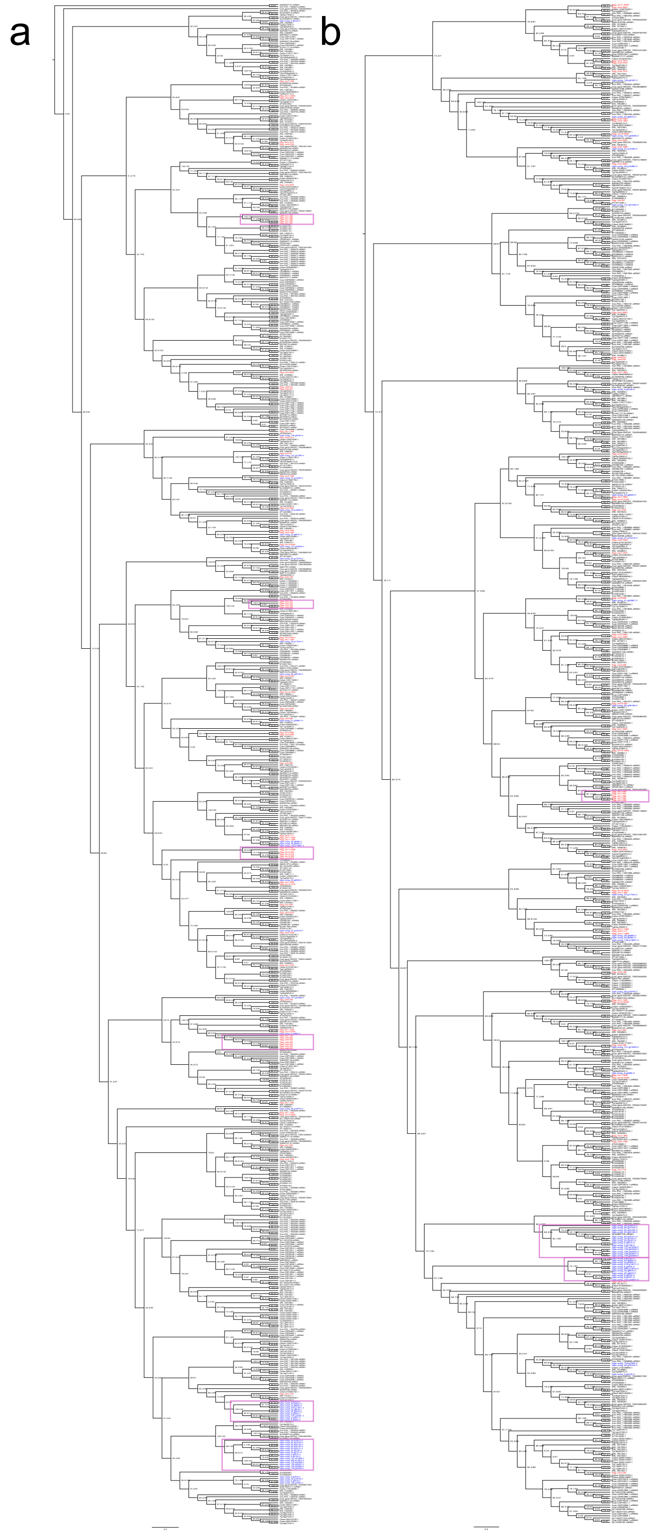

Figure S15: Peroxidase 10-like CDS and protein trees. (a) CDS sequences. (b) Protein sequences. All clades containing *Pinguicula gigantea* sequences with three or more tandem genes are highlighted and zoomed in. Branch support is the ultrafast bootstrap replicate score. *Utricularia gibba* sequences are in blue and *Pinguicula gigantea* sequences are in red. Tandem duplicates of 3 or more that group together on the tree are boxed in purple.

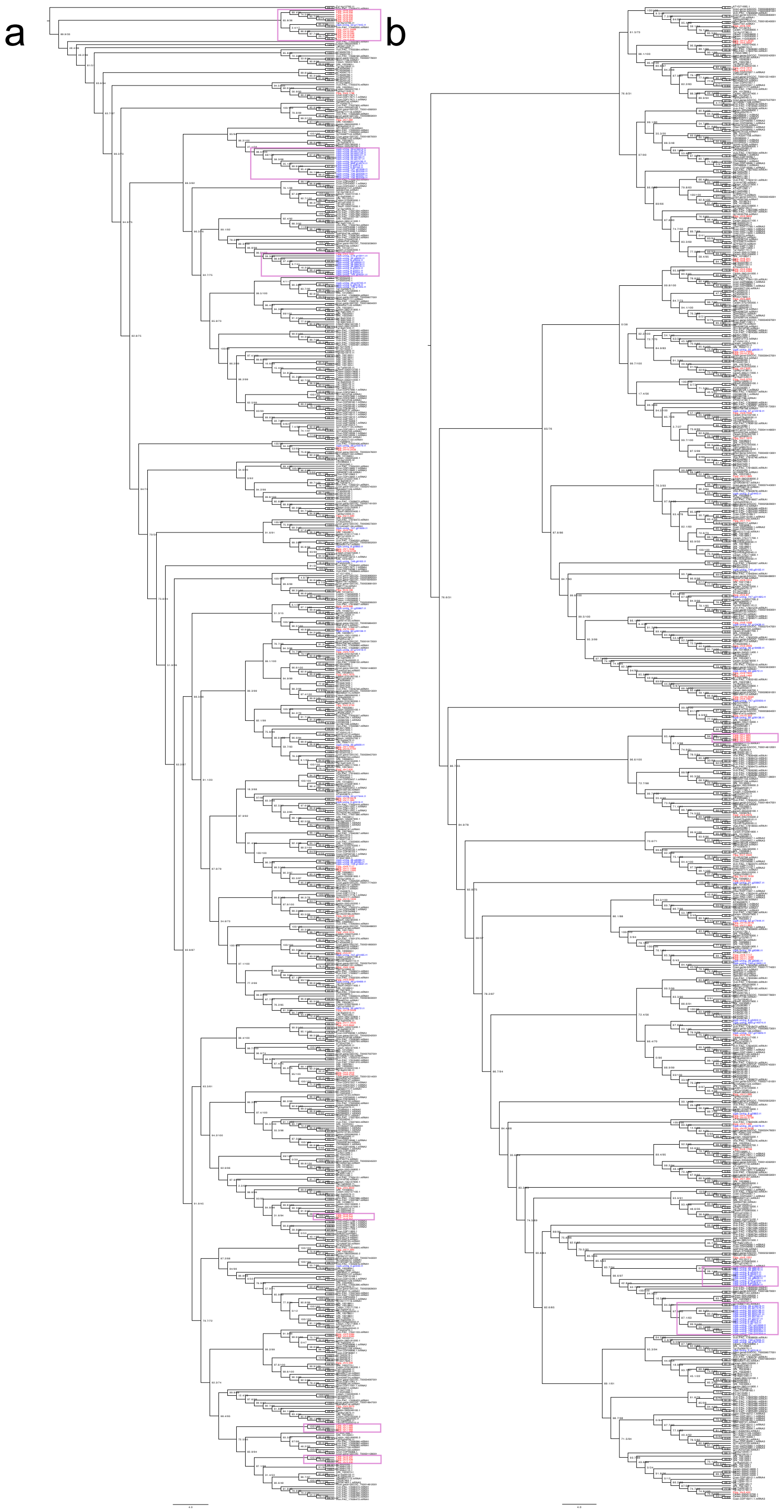


Figure S16: Putative peroxidase CDS and protein trees. (a) CDS sequences. (b) Protein sequences. All clades containing *Pinguicula gigantea* sequences with three or more tandem genes are highlighted and zoomed in. Branch support is the ultrafast bootstrap replicate score. *Utricularia gibba* sequences are in blue and *Pinguicula gigantea*

sequences are in red. Tandem duplicates of 3 or more that group together on the tree are boxed in purple.

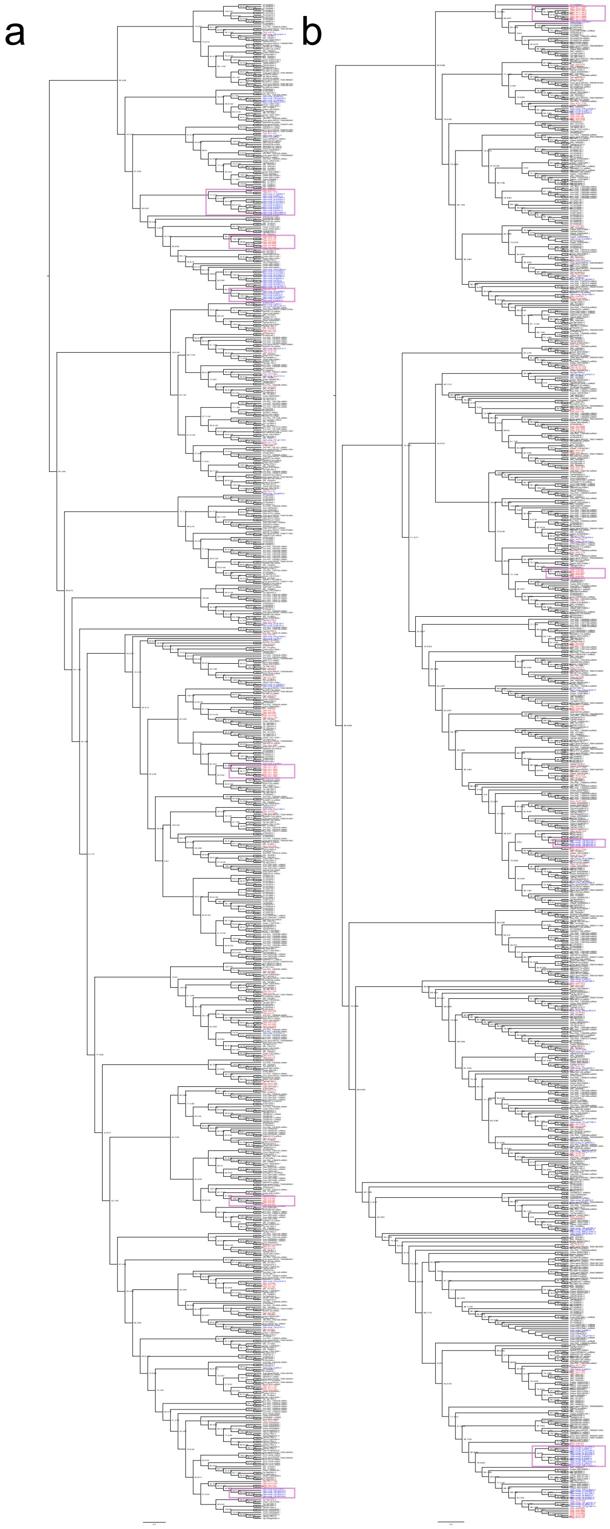


Figure S17: Predicted esterase/lipase CDS and protein trees. (a) CDS sequences. (b) Protein sequences. All clades containing *Pinguicula gigantea* sequences with three or more tandem genes are highlighted and zoomed in. Branch support is the ultrafast bootstrap replicate score. *Utricularia gibba* sequences are in blue and *Pinguicula gigantea* sequences are in red. Tandem duplicates of 3 or more that group together on the tree are boxed in purple.

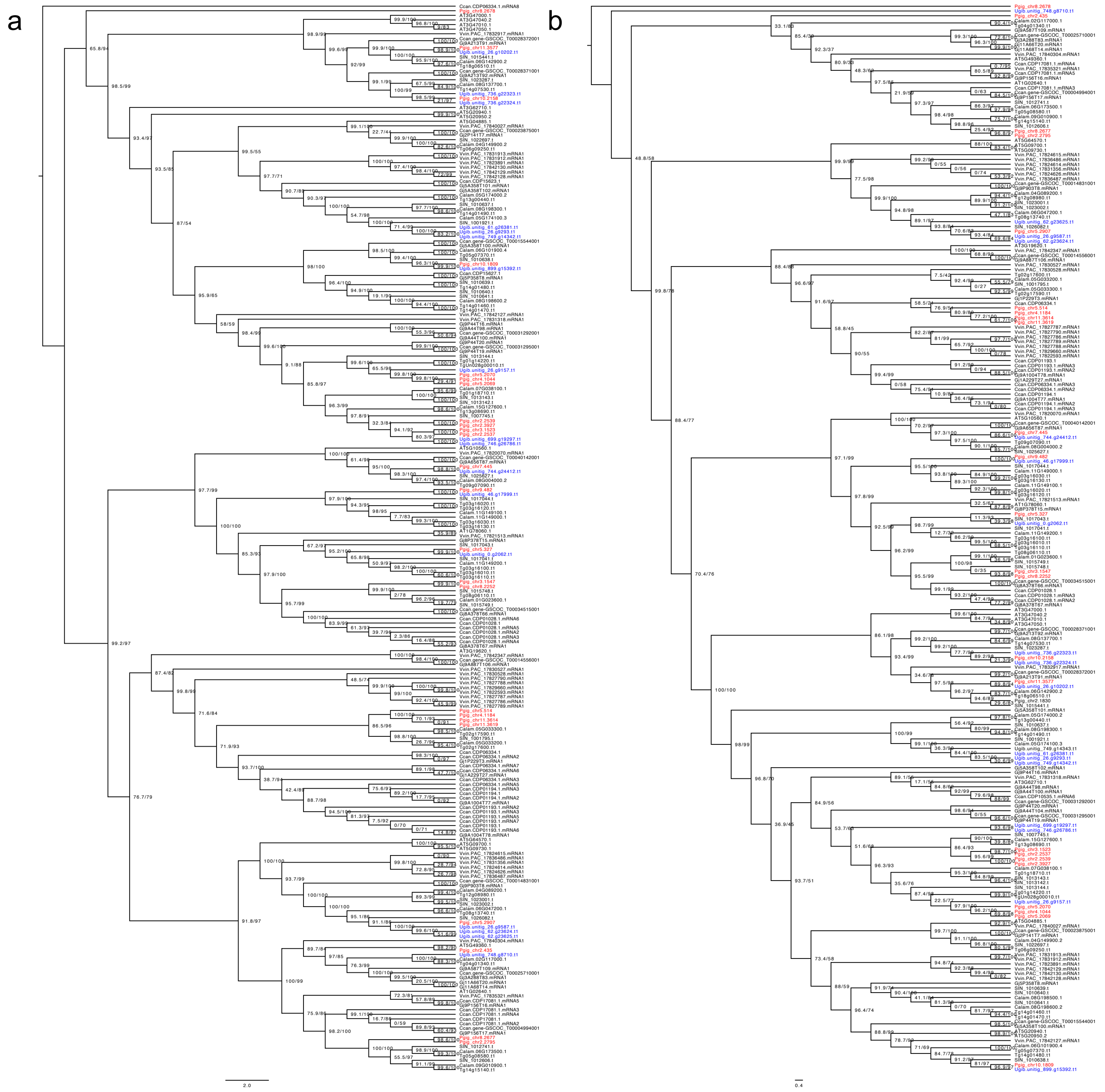


Figure S18: Putative beta-xylosidase CDS and protein trees. (a) CDS sequences. (b) Protein sequences. Branch support is the ultrafast bootstrap replicate score. *Utricularia gibba* sequences are in blue and *Pinguicula gigantea* sequences are in red. Tandem duplicates of 3 or more that group together on the tree are boxed in purple.

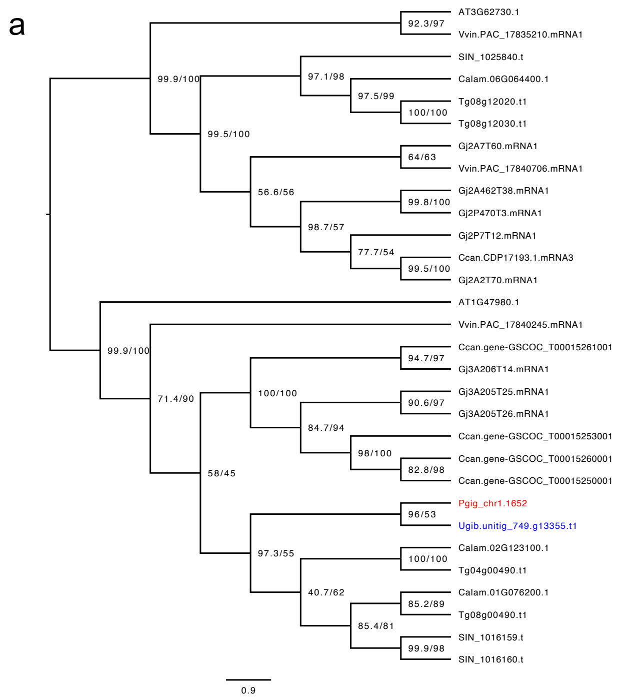

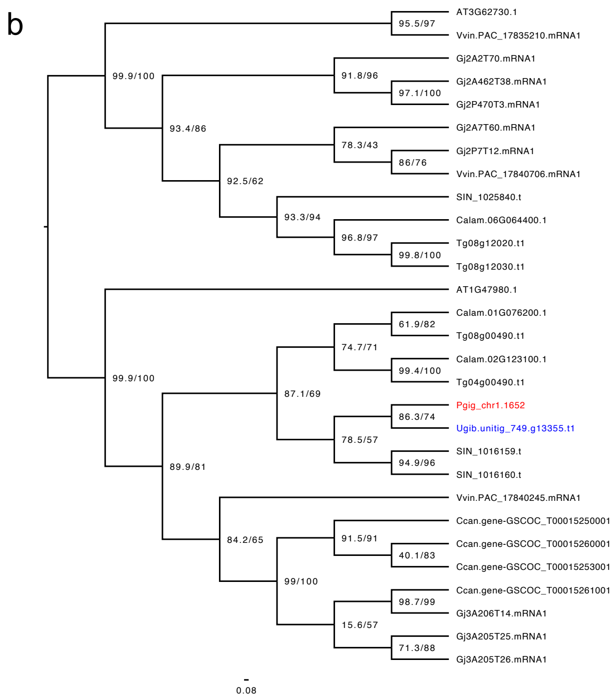


Figure S19: Desiccation-related protein CDS and protein trees. (a) CDS sequences. (b) Protein sequences. Branch support is the ultrafast bootstrap replicate score. *Utricularia gibba* sequences are in blue and *Pinguicula gigantea* sequences are in red. Tandem duplicates of 3 or more that group together on the tree are boxed in purple.

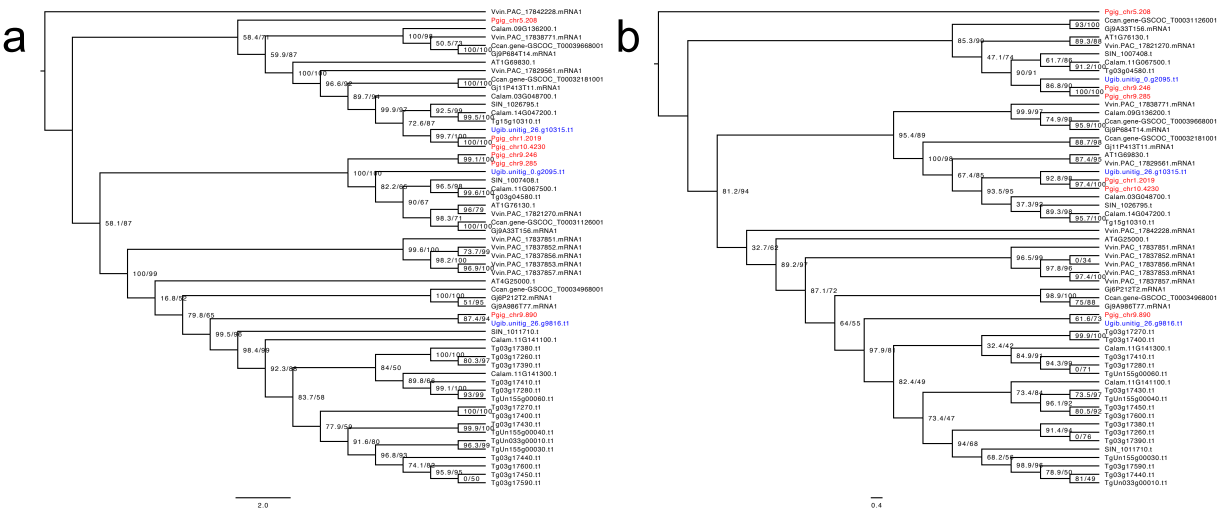


Figure S20: Alpha-amylase CDS and protein trees. (a) CDS sequences. (b) Protein sequences. Branch support is the ultrafast bootstrap replicate score. *Utricularia gibba* sequences are in blue and *Pinguicula gigantea* sequences are in red. Tandem duplicates of 3 or more that group together on the tree are boxed in purple.

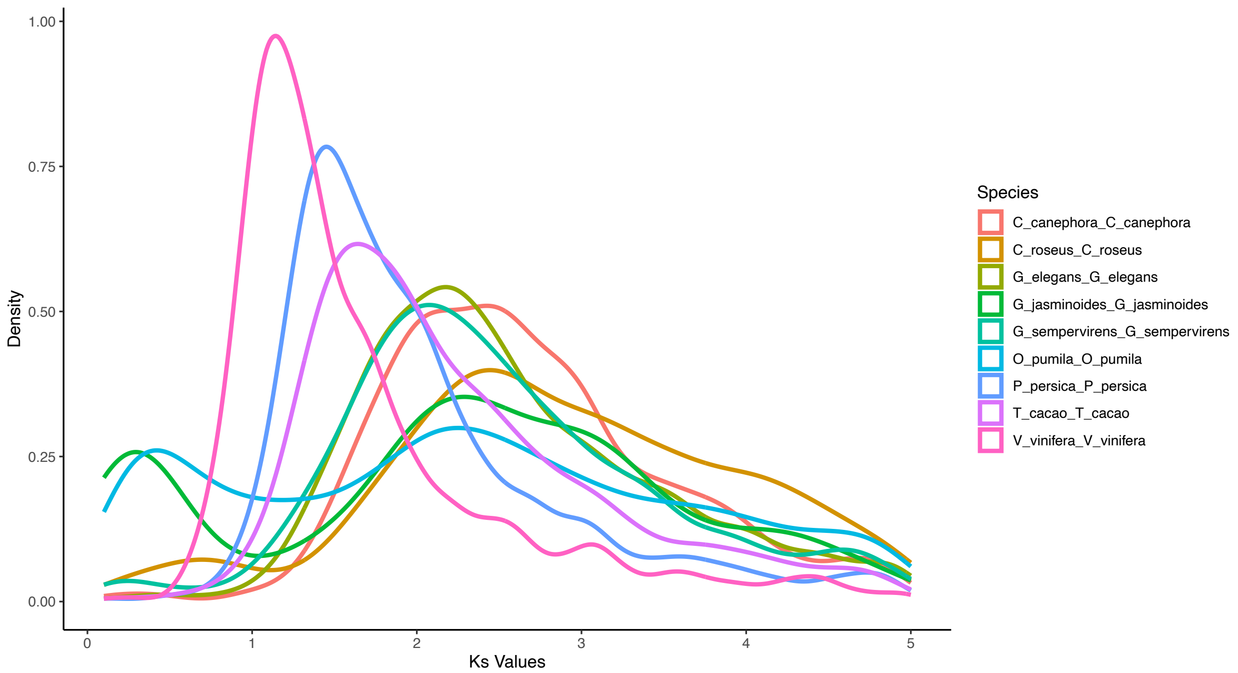


Figure S21: Density plots for Ks peaks representing the relative timing of the *gamma* hexaploidy event shared by all core eudicot lineages. Color key for Ks peak curved on right. The species included are as follows: *Coffea canephora*, *Catharanthus roseus*, *Gelsemium elegans*, *Gardenia jasminoides*, *Gelsemium sempervirens*, *Ophiorrhiza pumila*, *Prunus persica*, *Theobroma cacao*, and *Vitis vinifera*.
